# Functional balance between inhibitory and disinhibitory pathways predicts the heterogeneous effects of arousal on cortical timescales

**DOI:** 10.64898/2026.11.25.768094

**Authors:** Fabio Veneto, Luca Mazzucato, Julijana Gjorgjieva

**Affiliations:** School of Medicine and Health, Institute for Neuroscience, Technical University of Munich, Munich, Germany; School of Life Sciences, Technical University of Munich, Freising, Germany; Institute of Neuroscience, University of Oregon, Eugene, OR, USA; Munich Cluster for Systems Neurology (SyNergy), Munich, Germany

## Abstract

Cortical activity exhibits diverse timescales that vary with behavioral state, but the underlying circuit mechanisms remain unclear. Analyzing Allen Institute Neuropixels recordings from mouse primary visual cortex, we found that the fraction of excitatory neurons activated by optogenetic stimulation of somatostatin (SST) interneurons varied across mice. Intrinsic timescales lengthened during arousal in mice with a lower fraction of activated excitatory neurons and shortened in those with a higher fraction. To investigate a possible mechanism, we trained spiking networks with cell-type-specific plasticity, producing configurations dominated by direct SST inhibition of excitatory neurons or indirect disinhibition via parvalbumin interneurons. Arousal-like input to vasoactive intestinal peptide (VIP) interneurons slowed assembly dynamics, lengthened intrinsic timescales, and accelerated assembly formation when direct inhibition dominated, with opposite effects when indirect disinhibition dominated. Our results suggest that the balance between these pathways can account for opposite timescale responses to arousal and shape its computational consequences.

## Introduction

In the absence of external sensory inputs, the cortex generates structured spontaneous activity, which reflects learned structure and internal representations shaped by sensory experience and ongoing computations [1–6]. Neuronal fluctuations during spontaneous activity exhibit a wide range of intrinsic timescales [5, 7–9] that vary across cortical areas and individuals and define the windows over which circuits integrate information [7, 10–12]. Behavioral states such as wake-fulness, movements, and arousal rapidly modulate ongoing cortical activity, often on timescales faster than seconds [3, 13–22]. Yet, the circuit mechanisms that determine the direction and magnitude of state-dependent changes in intrinsic timescales remain unclear.

These state-dependent changes in cortical activity are mediated in part by modulatory and longrange signals that preferentially recruit specific interneuron classes [14–16, 23]. Three major interneuron classes expressing parvalbumin (PV), somatostatin (SST), and vasoactive intestinal peptide (VIP) account for more than 80% of GABAergic interneurons in the mouse cortex [24]. Together with excitatory neurons, these interneuron classes form a canonical cortical microcircuit whose subtype-specific connections support both direct inhibition and indirect disinhibition [25, 26]. These pathways provide multiple ways to regulate gain, temporal integration, and the balance between internally generated and stimulus-driven activity [24–28]. VIP interneurons, for example, are preferentially recruited during locomotion and arousal across visual, somatosensory, and auditory cortex [14–18, 29]. SST interneurons occupy a key branch point within the local circuit. They inhibit excitatory neurons directly, but can also inhibit PV interneurons and thereby indirectly disinhibit excitatory neurons [24–26]. Because VIP interneurons inhibit SST interneurons, their recruitment can disinhibit excitatory neurons by reducing direct SST-to-excitatory inhibition or inhibit them by releasing PV interneurons from SST inhibition. Even when the recruited interneuron class is similar, downstream effects on excitatory activity can differ across contexts, regions, and individuals, suggesting that differences in the relative influence of these pathways may lead to opposing responses to the same modulatory signal.

Previous models have shown how intrinsic cortical dynamics can be modulated by top-down inputs targeting either excitatory or inhibitory neurons [20, 30] and how VIP-mediated modulatory input can produce different responses depending on connectivity, network state, and synaptic dynamics [31, 32]. Related work has highlighted the importance of direct inhibitory and indirect disinhibitory pathways in shaping network stability, gain, and information processing [33–36]. In recurrent networks, spontaneous activity can switch between transiently active groups of strongly connected neurons, known as assemblies, giving rise to metastable dynamics [4, 20]. This raises the question of how the relative influence of direct inhibitory and indirect disinhibitory pathways shapes metastable activity and state-dependent intrinsic timescales in networks whose connectivity develops through synaptic plasticity. These differences in circuit organization may also have consequences for the formation of new assemblies.

Here, we combined publicly available *in vivo* recordings and optogenetic stimulation data from the Allen Institute ‘Visual Coding’ Neuropixels dataset [37] with computational modeling to investigate how inhibitory circuit organization shapes state-dependent dynamics. We first found substantial variation across mice in the excitatory responses to SST activation. To investigate a possible circuit mechanism, we trained recurrent spiking networks to form excitatory assemblies. In one configuration, SST-to-excitatory synapses strengthened through local plasticity, favoring direct inhibition; in the other, these synapses remained fixed at their initial weak values, favoring indirect disinhibition through PV interneurons. Although the resulting network configurations exhibited similar spontaneous activity, the same SST modulatory input accelerated assembly dynamics in the inhibition-dominated circuit but slowed them down in the disinhibition-dominated configuration. By combining the two SST pathways within a single network, we reproduced the continuum of excitatory responses to SST activation observed across mice. In the experimental recordings, we found that the fraction of excitatory neurons activated by SST stimulation predicted the direction of arousal-dependent changes in intrinsic timescales. Applying arousal-like VIP modulation generated opposite effects on the formation of new assemblies in the two configurations consistent with the changes in intrinsic timescales. Together, our findings suggest that similar spontaneous activity can arise from different inhibitory circuit configurations, with distinct consequences for state-dependent dynamics and the formation of new assemblies.

## Results

### Excitatory responses to SST interneuron activation vary across individual mice *in vivo*

Top-down and behavioral-state signals recruit distinct classes of cortical interneurons (INs), but their effects on excitatory activity depend on behavioral context and the local circuit in which these interneurons are embedded [14–16, 23, 25]. To examine whether the consequences of recruiting a single interneuron class also vary across animals, we analyzed the Allen Institute ‘Visual Coding’ Neuropixels dataset [37], which combines large-scale recordings across the mouse visual hierarchy during behavior with optogenetic activation of genetically defined interneuron populations (Fig. 1A). We focused on SST INs in the primary visual cortex (V1) because their recruitment can have opposing effects on excitatory activity through direct inhibition and indirect disinhibition via PV INs. We reasoned that the proportions of suppressed and activated excitatory neurons following SST activation could provide a functional indication of the relative influence of these two pathways. SST INs are also prominent targets of neuromodulatory and long-range inputs. For instance, cholinergic input to V1 directly facilitates SST INs [38], locomotion recruits them in S1 [17], transitions to high arousal increase their activity in posterior parietal cortex (PPC) [39], and cortico-cortical feedback from the lateromedial area (LM) to V1 preferentially targets them [40].

**Fig. 1.**
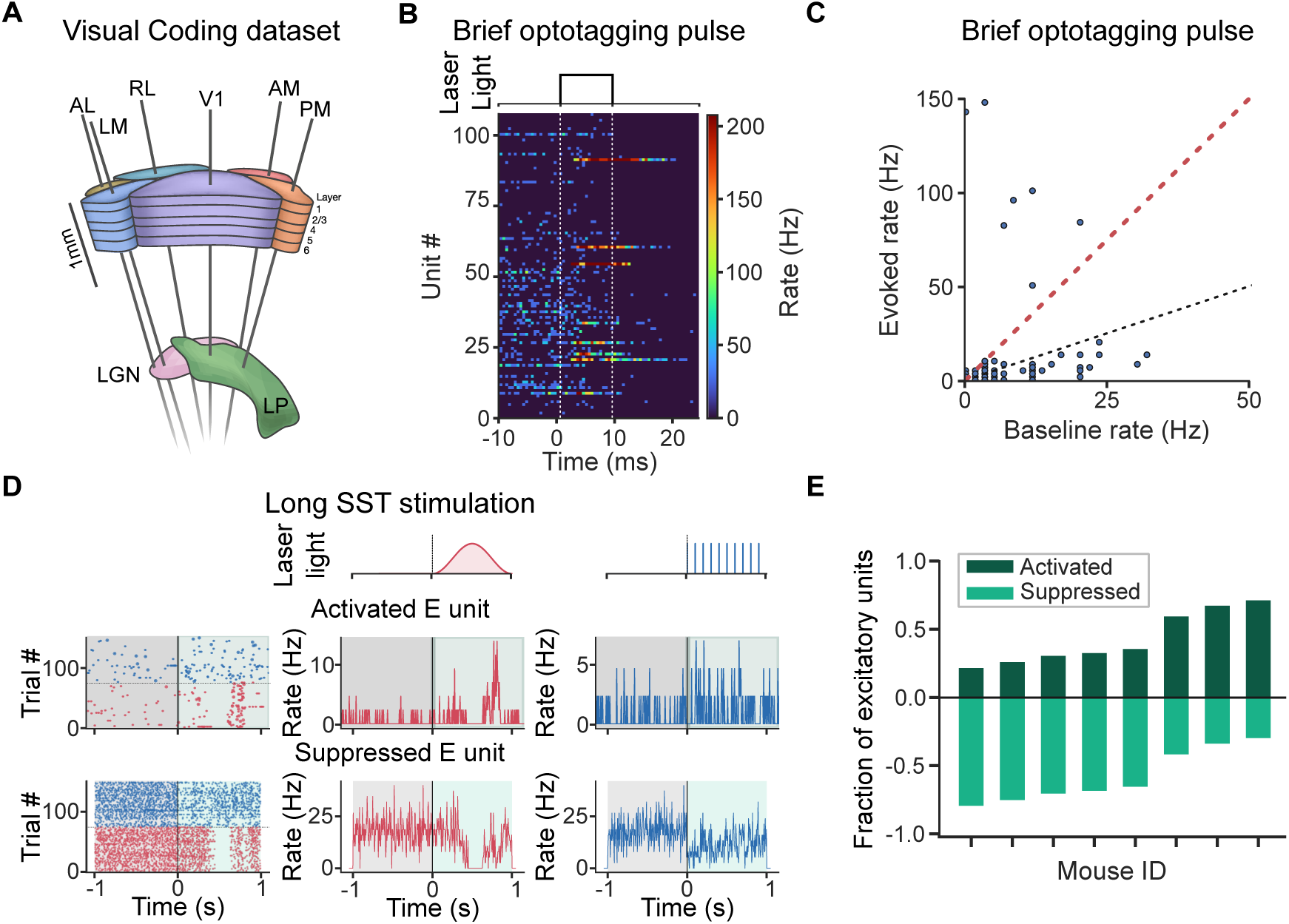
A continuum of excitatory responses to optogenetic SST activation in V1. **A.** Schematic of the mouse visual system, showing the six cortical visual areas (V1, LM, RL, AL, PM, AM) and the thalamic nuclei LGN and LP, recorded simultaneously with Neuropixels probes in the Allen Institute ’Visual Coding’ dataset. **B.** Optotagging of SST INs in one example *Sst* -Cre session. (Top) Optogenetic light stimulus consisting of a single 10 ms pulse. (Bottom) Firing rate of every V1 unit aligned to pulse onset; vertical white dashed lines mark pulse onset and offset. **C.** Evoked versus baseline firing rate for every unit in the same session (evoked window +1 to +9 ms, baseline window *−*10 to *−*2 ms relative to pulse onset). Units above the red dashed threshold were classified as optotagged SST INs (see Methods); the black dotted line is the identity. **D.** Responses of two example V1 excitatory (E) units to *∼* 1 s optogenetic activation of SST INs, delivered as either a raised-cosine pulse (red) or a train of fast pulses (blue). The light waveforms are shown above. For each unit, single-trial spike rasters for both stimulation protocols are shown on the left, followed by the trial-averaged firing rates during the raised-cosine pulse (middle, red) and the train of fast pulses (right, blue). The top row shows an E unit activated by SST drive, and the bottom row an E unit suppressed by SST drive. Gray shading marks the pre-stimulus baseline window, while the colored post-onset shading marks the stimulation period (see Methods). **E.** Fraction of V1 E units activated (dark green, plotted upward) or suppressed (light green, plotted downward) by SST drive in each mouse. Fractions are normalized by the number of E units with a nonzero firing rate change.

We first identified optotagged SST INs by analyzing brief (10 ms) light pulses in *Sst* -Cre mice expressing channelrhodopsin-2 (ChR2) in SST INs (Fig. 1B). We labeled SST units as those whose firing rate during the light pulse exceeded three times their baseline rate before the light pulse (Fig. 1B,C; see Methods). We then used longer (*∼* 1 s) optogenetic activation of SST INs, delivered either as a raised-cosine pulse or as a train of fast pulses, to determine how this perturbation affected excitatory neurons. For each excitatory (E) unit, we compared its firing rate immediately before and during the stimulus (see Methods). Both response types coexisted within individual V1 recordings: some E units were suppressed by SST activation, consistent with direct SST-to-E inhibition, while others were activated, consistent with PV-mediated indirect disinhibition (Fig. 1D).

The balance between the two response types varied widely across individual mice (Fig. 1E). In most mice, the majority of E units were suppressed by SST drive, consistent with a stronger relative contribution of direct SST-to-E inhibition. In a few mice, however, the majority of E units were activated, consistent with a stronger relative contribution of indirect disinhibition. For each mouse, we summarized this balance by the fraction of SST-activated E units, frac_up_ (see Methods), which we use below as an experimentally accessible measure of the net effect of SST activation on the local excitatory population in V1. As a control, we examined excitatory responses to opto-genetic stimulation in five *Pvalb*-Cre mice expressing ChR2 selectively in PV INs. In contrast to SST activation, PV activation did not activate more E units than it suppressed in any of the five mice (Fig. S1D). The relative prevalence of activated and suppressed E units also differed across cortical layers (Fig. S1A; see Methods). These observations reveal a functional continuum across mice, ranging from predominantly suppressive to predominantly activating excitatory responses to SST drive. We hypothesized that a variation in the functional balance between direct and indirect SST pathways could generate this response continuum and examined what consequences it would have for state-dependent cortical dynamics.

### Shaping inhibitory and disinhibitory pathways by training network assemblies with cell-type specific plasticity

To understand how the same SST drive can suppress excitatory neurons in one mouse and activate them in another, and what this implies for state-dependent dynamics, we built spiking network models containing excitatory neurons and the PV, SST, and VIP interneuron classes introduced above. The populations were connected according to the canonical cortical microcircuit, including reciprocal connections between excitatory and PV INs, projections from SST INs to excitatory and PV INs, and reciprocal connections between SST and VIP INs [23–25] (Fig. 2A). This architecture incorporates the direct SST-to-excitatory and indirect SST-to-PV-to-excitatory pathways described above.

**Fig. 2.**
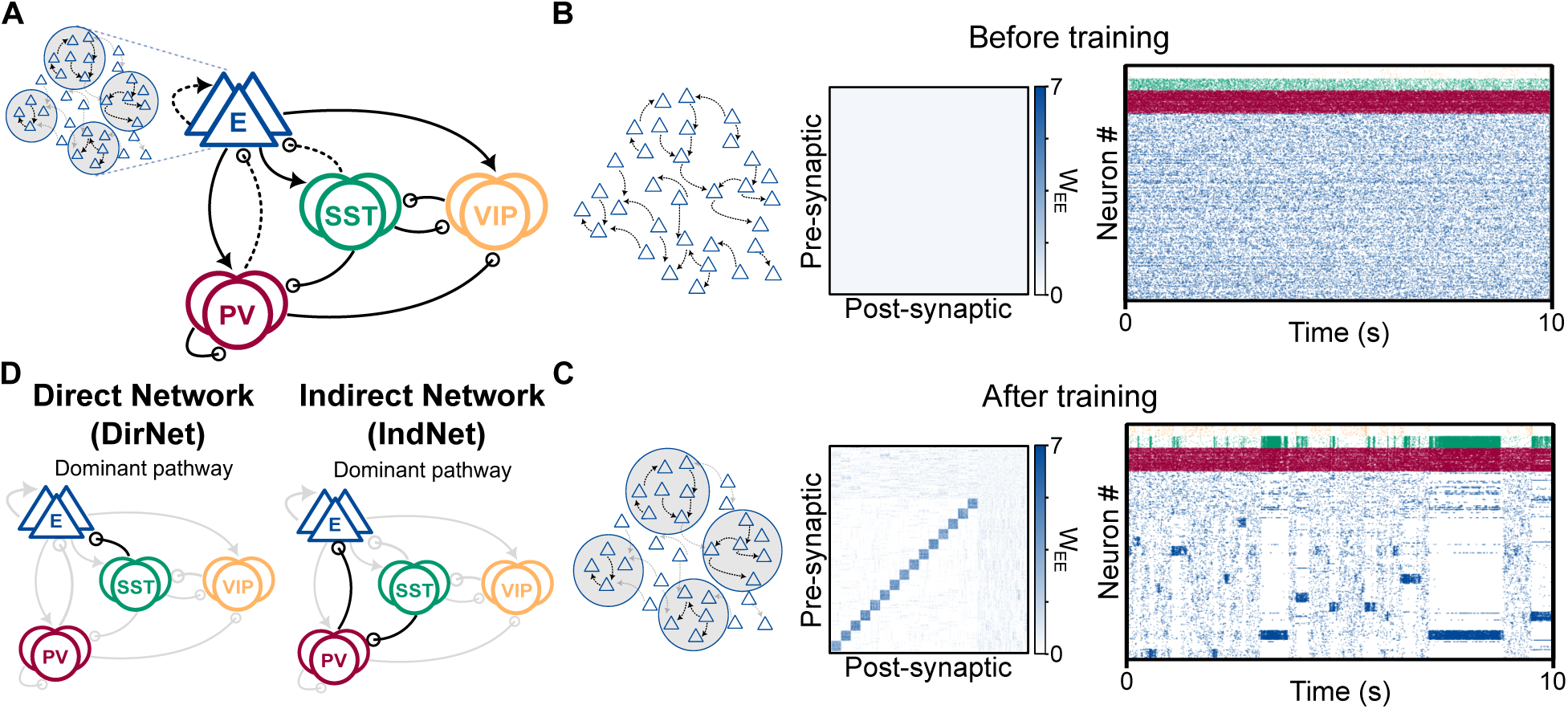
Cell-type-specific plasticity generates excitatory assemblies with distinct inhibitory connectivity. **A.** Schematic of the clustered spiking neural network with multiple interneuron types. Plastic connections include E-to-E, PV-to-E, and SST-to-E. In DirNet, SST-to-E synapses are plastic, while in IndNet they are static and weak. **B.** Network before training, illustrated for DirNet. From left to right: schematic of homogeneous excitatory connectivity, excitatory connectivity matrix, and raster plot showing excitatory (blue), PV (red), SST (green) and VIP (orange) spikes over 10 s. **C.** Same as **B** after training, showing the formation of excitatory assemblies and spontaneous transitions between assembly-specific patterns of activity. **D.** Dominant pathways from SST to excitatory neurons in DirNet (left) and IndNet (right). In DirNet, SST-to-E synapses strengthen during training, and the direct pathway dominates. In IndNet, these synapses remain weak, and SST INs influence excitatory neurons mainly through PV INs.

Experimental studies have revealed that excitatory neurons are further organized into non-random subnetworks, with more probable and stronger connectivity between neurons that share similar stimulus preferences, such as selectivity for the same orientation of a visual stimulus [2, 41–44]. These interconnected groups of neurons are commonly referred to as assemblies [45]. Such structured connectivity is thought to emerge through experience-dependent plasticity, which refines synaptic strengths based on patterned neuronal activity [46–48].

We initialized the networks with homogeneous connectivity, resulting in asynchronous irregular activity (Fig. 2B, Fig. S2). To model the emergence of structured connectivity during development, we implemented an unsupervised learning protocol using biologically plausible experience-dependent plasticity rules. We exposed the excitatory population to correlated input and trained the synaptic connections between excitatory neurons using a triplet spike-timing-dependent plasticity (STDP) rule, known to form assemblies [49–51] (Fig. 2A-C, Fig. S3, Fig. S2, see Methods). For inhibitory synapses onto excitatory neurons, we implemented cell-type-specific inhibitory STDP (iSTDP). PV-to-excitatory synapses followed an established iSTDP rule that stabilizes excitatory firing rates [52–54]. SST-to-excitatory synapses followed a distinct iSTDP rule with a different temporal kernel, based on recent experimental measurements of SST-specific inhibitory plasticity [54].

To vary the relative influence of the direct and indirect SST pathways, we considered two network configurations with and without SST-to-excitatory plasticity: the ‘Direct Network’ (*DirNet*) and the ‘Indirect Network’ (*IndNet*). In DirNet, SST-to-E synapses strengthened rapidly through plasticity during training, such that SST INs influenced excitatory neurons predominantly through the direct SST-to-E pathway (Fig. 2D, Fig. S3). In IndNet, SST-to-E plasticity was absent, leaving these synapses much weaker especially after training. Consequently, SST INs influenced excitatory neurons mainly through the indirect SST-to-PV-to-excitatory pathway (Fig. 2D, Fig. S3).

In both network configurations, training strengthened connections within groups of excitatory neurons while connections between groups remained weak, thereby generating excitatory assemblies (Fig. 2C, Fig. S3, Fig. S2). After training, excitatory neurons exhibited structured spontaneous activity even in the absence of sensory input. This activity was organized into intermittent periods of strong co-activation within individual assemblies. At any given time, one assembly tended to dominate while others were largely silent (Fig. 2C, Fig. S2). The network thus transitioned irregularly between discrete, assembly-specific patterns of activity known as metastable states [4, 20, 22, 30, 50, 55]. These patterns of spontaneous activity resembled those observed experimentally [3, 4, 56].

Despite their differences in SST-to-excitatory connectivity, DirNet and IndNet exhibited similar baseline firing activity and metastable dynamics (Fig. 2C, Fig. S2). The two network configurations therefore could not be distinguished based on their spontaneous activity. However, because the direct and indirect SST pathways have opposite net effects on excitatory neurons, the two configurations generate a clear prediction: the same top-down input to SST INs should suppress excitatory activity in DirNet but enhance it in IndNet. DirNet and IndNet therefore allowed us to ask whether differences in the relative influence of the direct and indirect SST pathways could account for the predominantly suppressive and predominantly activating responses observed across mice in Fig. 1E. We next tested this prediction by directly stimulating SST INs in each network.

### The balance between direct and indirect SST pathways shapes intrinsic timescales

We first asked whether the two network configurations reproduced the predominantly suppressive or activating excitatory responses to SST activation observed *in vivo* (Fig. 1E). To mimic the ex-perimental perturbation, we applied an additional input current to the SST population after training (see Methods). In DirNet, SST drive strongly suppressed excitatory firing (Fig. 3A,E), consistent with mice in which most excitatory units were suppressed by SST activation. In IndNet, SST drive suppressed PV interneurons and thereby increased excitatory firing (Fig. 3B,G), consistent with mice in which most excitatory units were activated by SST activation. DirNet and IndNet therefore reproduced the two ends of the functional response continuum observed across mice.

**Fig. 3.**
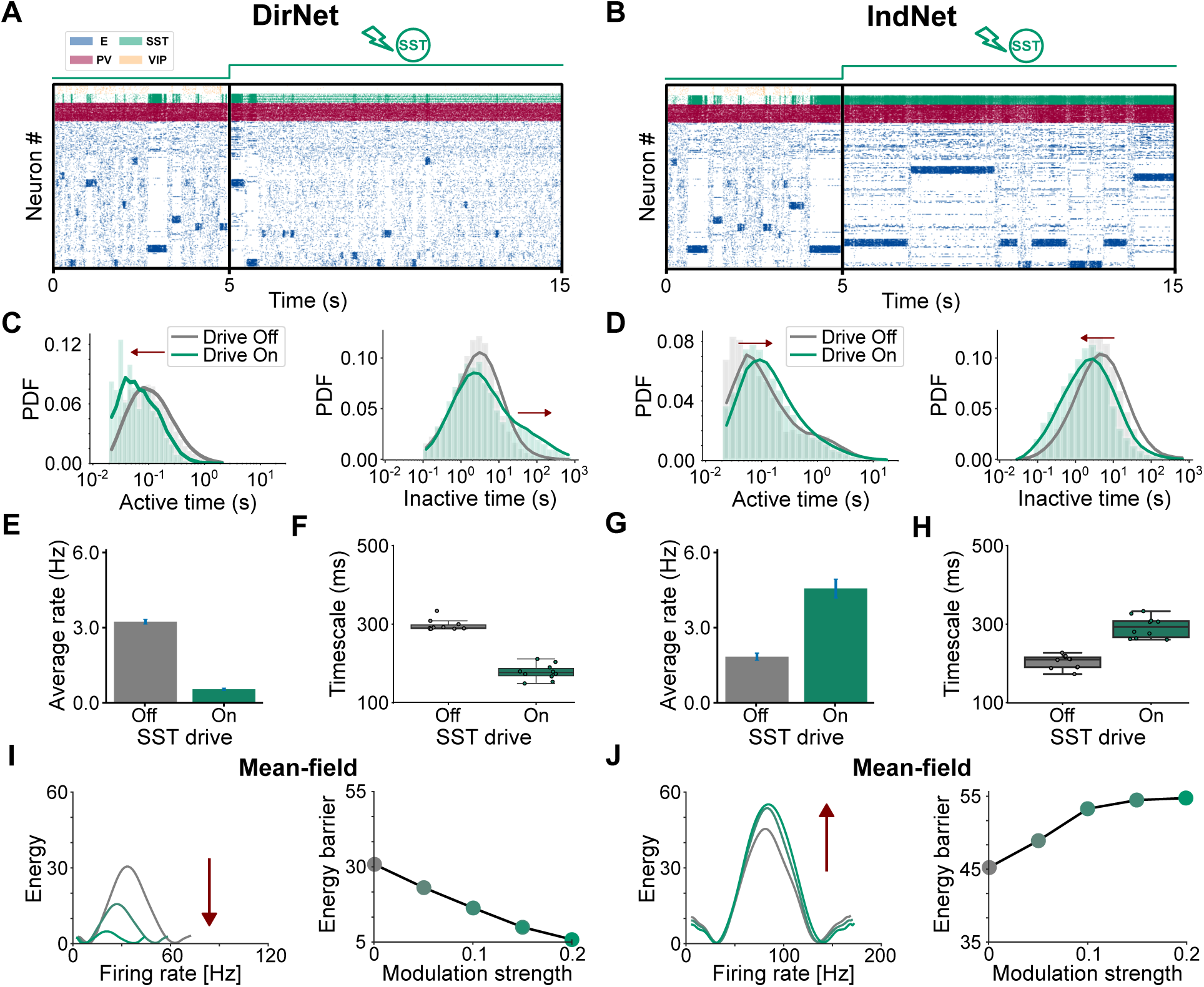
SST drive accelerates or slows assembly dynamics depending on the dominant pathway. **A.** Raster plot of DirNet activity without SST drive (first 5 s) and with SST drive (last 10 s), showing shorter assembly activations during SST drive. **B.** Same as **A** for IndNet. SST drive results in longer assembly activations. **C.** Probability density functions (PDFs) of active and inactive times in DirNet. Active times measure how long an assembly remains continuously active, whereas inactive times measure the interval between two consecutive activations of the same assembly (continuous asynchronous irregular activity for that assembly). Conditions with (green) and without (gray) SST drive are shown. **D.** Same as **C** for IndNet. **E.** Average firing rate of excitatory neurons in DirNet with (green) and without (gray) SST drive. **F.** Intrinsic timescales of excitatory neurons in DirNet with (green) and without (gray) SST drive. **G,H.** Same as **E,F** for IndNet. **I.** (Left) Estimate of a one-dimensional energy landscape for a single assembly as a function of assembly firing rate. Low firing rate corresponds to asynchronous irregular activity, while high firing rate corresponds to a single assembly being continuously active. (Right) Height of the energy barrier separating the low- and high-firing states as a function of SST modulation strength. **J.** Same as **I** for IndNet.

We then hypothesized that top-down modulation of SST could modulate the network metastable activity in opposite directions in DirNet and IndNet. We captured the metastable dynamics of transient assembly activation in terms of the time an assembly was continuously active as its ‘active time’, and the time between two consecutive activations of the same assembly (resulting in a continuous asynchronous irregular state for an assembly) as the ‘inactive time’ (see Methods). We found that, in DirNet, SST drive accelerated assembly dynamics (Fig. 3A): active times shortened while inactive times lengthened (Fig. 3C). In contrast, in IndNet, SST drive slowed assembly dynamics (Fig. 3B): active times lengthened while inactive times shortened (Fig. 3D).

To link these changes with a quantity that can be measured *in vivo*, we computed the intrinsic timescale of each neuron from the temporal autocorrelation of its activity, with and without top-down modulation of SST [7] (see Methods). Consistent with the shorter assembly activations, SST drive shortened the intrinsic timescale in DirNet, indicating that activity decorrelated more rapidly over time (Fig. 3F). Consistent with the longer assembly activations, SST drive lengthened the intrinsic timescale in IndNet (Fig. 3H).

To identify the dynamical principle underlying these opposite changes in assembly dynamics, we obtained a mean-field approximation of both DirNet and IndNet network configurations, in which top-down modulation shifted the mean input current to the SST population (see Methods). Within an attractor landscape framework obtained from the mean field theory, the low-firing asynchronous state and the high-firing assembly-active state correspond to two stable states separated by an energy barrier (Fig. 3I,J; see Methods) [20, 22, 30]. In DirNet, increasing the strength of SST modulation lowered the energy barrier, making transitions between states more frequent and shortening active times (Fig. 3I). In IndNet, it raised the barrier, making transitions rarer and prolonging active times (Fig. 3J).

In sum, our results show that the same top-down input to SST INs can drive activity in opposite directions, depending on whether the network architecture is DirNet or IndNet. This flexibility captures the suppressive and activating ends of the response continuum observed across mice. In the model, the relative influence of the DirNet vs. IndNet SST pathways determines both the sign of the excitatory response and whether SST drive accelerates or slows assembly dynamics and shortens or lengthens intrinsic timescales.

### Bidirectional modulation of intrinsic timescales by arousal-like VIP input

Behavioral-state signals recruit multiple interneuron classes. In V1, VIP INs depolarize during pupil dilation, even during quiet wakefulness [29]. In turn, VIP INs inhibit SST INs [24, 25]. We therefore modeled arousal-like modulation as an additional input current to the VIP population and asked how the two network configurations responded (Fig. 4A,B). In DirNet, VIP activation suppressed SST INs, thus reducing SST-mediated inhibition of excitatory neurons, whose active times and average firing rate increased (Fig. 4C,E). In contrast, in IndNet, VIP activation suppressed SST INs and thereby disinhibited PV INs, increasing PV-mediated inhibition of excitatory neurons, whose active times and average firing rate decreased (Fig. 4D,G). Inactive times changed only slightly in both network configurations (Fig. 4C,D). Accordingly, VIP drive lengthened the intrinsic timescale in DirNet, consistent with the longer assembly active times (Fig. 4F), and shortened it in IndNet, consistent with the shorter assembly active times (Fig. 4H).

**Fig. 4.**
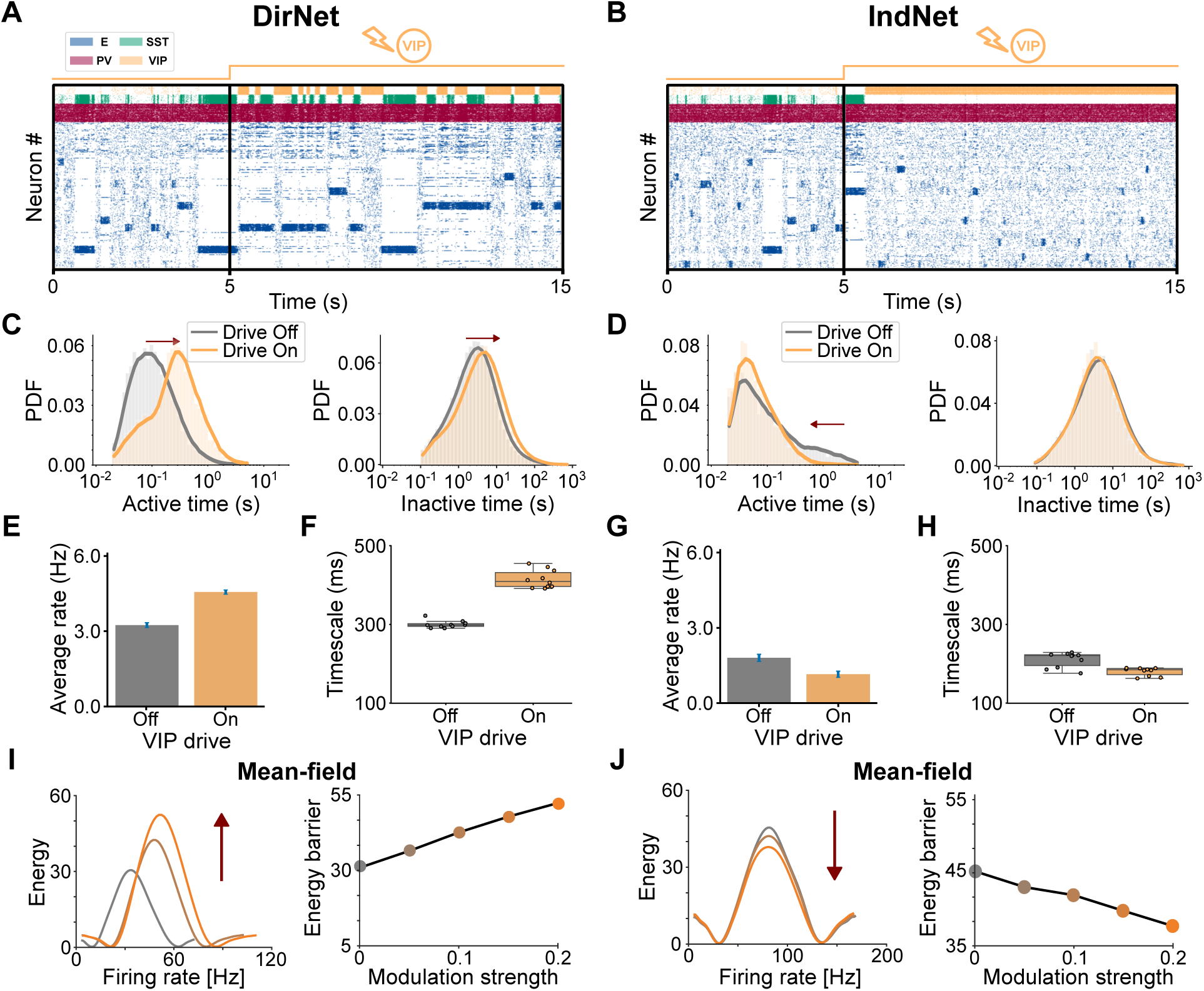
VIP drive reverses the effects of SST modulation on network dynamics. **A.** Raster plot of DirNet activity without VIP drive (first 5 s) and with VIP drive (last 10 s), showing longer assembly activations during VIP drive. **B.** Same as **A** for IndNet. VIP drive results in shorter assembly activations. **C.** Probability density functions (PDFs) of active and inactive times in DirNet, defined as in Fig. 3C. Conditions with (orange) and without (gray) VIP drive are shown. **D.** Same as **C** for IndNet. **E.** Average firing rate of excitatory neurons in DirNet with (orange) and without (gray) VIP drive. **F.** Intrinsic timescales of excitatory neurons in DirNet with (orange) and without (gray) VIP drive. **G,H.** Same as **E,F** for IndNet. **I.** (Left) Estimate of a one-dimensional energy landscape for a single assembly as a function of assembly firing rate. Low firing rate corresponds to asynchronous irregular activity, while high firing rate corresponds to a single assembly being continuously active. (Right) Height of the energy barrier separating the low- and high-firing states as a function of VIP modulation strength. **J.** Same as **I** for IndNet.

The mean-field analysis showed a corresponding reversal of the changes in the energy barrier induced by SST drive. In DirNet, increasing VIP modulation strength raised the energy barrier, making transitions between states rarer and prolonging active times, whereas in IndNet it lowered the barrier, making transitions more frequent and shortening active times (Fig. 4I,J).

Taken together, these results show that VIP drive induces effects opposite of direct SST drive in each network configuration. Across both perturbations, the direction of change in assembly dynamics and intrinsic timescales follows the net effect on excitatory neurons: disinhibition slows assembly dynamics and lengthens intrinsic timescales, whereas inhibition accelerates assembly dynamics and shortens intrinsic timescales. In particular, arousal-like input to VIP INs is predicted to lengthen intrinsic timescales in DirNet and to shorten them in IndNet.

### A mixed SST population supports flexible control of network dynamics

DirNet and IndNet represent network configurations in which either the direct or the indirect SST pathway dominates across the SST population. However, SST INs are highly diverse, comprising molecularly and morphologically distinct subtypes, such as Martinotti and non-Martinotti cells, that form cell-type-specific circuits [57] and are differentially regulated during learning [58, 59]. Functionally distinct SST subpopulations can also receive and gate specific long-range inputs, with only a subset of layer 2/3 SST INs receiving pulvinar input to V1 [60]. We therefore asked whether direct and indirect SST functional motifs could coexist within a single circuit and be recruited selectively.

To model the developmental trajectories of heterogeneous SST populations, we trained both connectivity motifs into a single mixed network. Specifically, we divided the SST population into two subgroups during training: in one subgroup SST-to-excitatory synapses were plastic, while in the other they were static and remained weak (Fig. 5A). After training, this resulted in one SST sub-group forming strong (DirNet-like) connections and another forming weak (IndNet-like) connections onto the excitatory population. Each excitatory neuron received a combination of strong and weak SST inputs, with the balance between the two determined by the proportion of neurons in each SST subgroup. The training protocol was otherwise the same as in DirNet and IndNet (Fig. S4, see Methods).

**Fig. 5.**
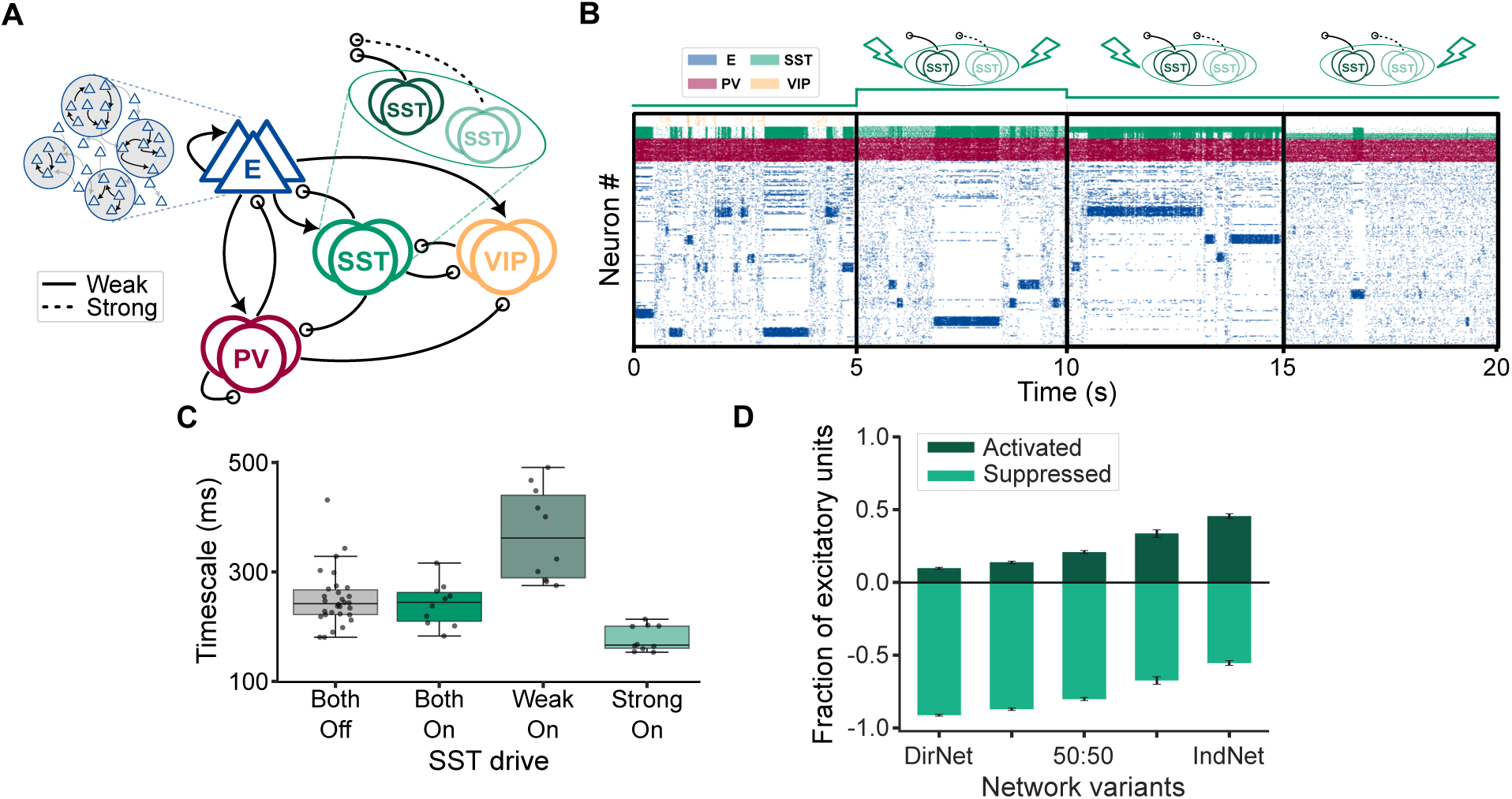
SST population composition and recruitment shape excitatory responses and intrinsic timescales. **A.** Schematic of the spiking recurrent network model. During training, excitatory-to-excitatory and PV-to-excitatory connections are plastic, while the SST population is divided into two subgroups: one subgroup experiences SST-to-excitatory plasticity producing strong connections (DirNet-like), while connections are static and weak in the other subgroup (IndNet-like). **B.** Raster plot of network activity without SST drive (before 5 s) and during top-down drive to both SST subgroups (between 5 s and 10 s), only the weakly projecting SST subgroup (between 10 s and 15 s), and only the strongly projecting SST subgroup (between 15 s and 20 s). **C.** Intrinsic timescales across trials without SST drive (gray) and during activation of both SST subgroups (green), only the weakly projecting subgroup (dark green) or only the strongly projecting subgroup (light green). **D.** Fractions of E units activated (plotted upward, dark green) or suppressed (plotted downward, light green) by SST drive in networks with different proportions of strongly and weakly projecting SST INs (DirNet 100:0, 75:25, 50:50, 25:75, IndNet 0:100), normalized to the total number of E units with a nonzero firing rate change. Error bars represent standard error of the mean (SEM) across multiple simulations.

After training, we activated either the full SST population, only the weakly projecting subgroup, or only the strongly projecting subgroup (Fig. 5B). When both subgroups were activated together, their opposing effects approximately canceled, leaving the intrinsic timescale unchanged (Fig. 5C). Selective recruitment of the two subgroups changed the intrinsic timescale in opposite directions: driving only the strong (DirNet-like) subgroup shortened the intrinsic timescale, while driving only the weak (IndNet-like) subgroup lengthened it. Thus, selective recruitment of SST subpopulations allowed the same network to shift between faster and slower intrinsic dynamics without changing its connectivity.

We next interpolated between IndNet and DirNet by varying the fraction of SST INs whose connections onto excitatory neurons were allowed to strengthen during training. For each network, we computed frac_up_, the fraction of excitatory neurons activated by SST drive, using the same measure as in the experimental data (Fig. 1E; see Methods). Increasing the proportion of strongly projecting SST INs shifted excitatory responses toward suppression (Fig. 5D). This suggests that differences in the distribution of SST-to-excitatory synaptic strengths could contribute to the variation in response balance observed across mice (Fig. 1E). Together, these results show that heterogeneity in SST-to-excitatory connectivity is sufficient to reproduce the continuum of excitatory responses across mice while allowing selective recruitment of SST subpopulations to bidirectionally modulate intrinsic timescales within a single network.

### Excitatory responses to SST activation predict arousal-dependent timescale changes across mice

Intrinsic timescales are strikingly heterogeneous *in vivo*, varying across cortical areas, layers, and individuals [6, 11, 12, 21]. While this variability is well documented, its circuit origin remains unclear. Our model predicts that the direction of arousal-dependent timescale modulation should covary with the excitatory response to SST activation. We tested this prediction in the same mice analyzed in Figure 1, for which frac_up_, the fraction of excitatory units activated by SST stimulation, was proposed as a functional readout of the relative influence of the two SST pathways.

We identified bouts of high and low arousal in each session from pupil diameter during the spontaneous condition (Fig. 6A, top; see Methods), and estimated the population intrinsic timescale of V1 excitatory units in sliding windows across the session (Fig. 6A, bottom; see Methods). For each mouse, we then computed the change in intrinsic timescale between high- and low-arousal bouts, Δ*τ* = *τ*_high_ *− τ*_low_. Δ*τ* varied markedly across individual mice, and its sign was predicted by the fraction of SST-activated units measured independently during optogenetic stimulation (Fig. 6B; Pearson *r* = *−*0.811, *p* = 0.015; Spearman *ρ* = *−*0.786, *p* = 0.021; *n* = 8 mice). Mice with low frac_up_, in which SST activation predominantly suppressed excitatory units (functionally DirNet-like), showed longer intrinsic timescales during high than during low arousal. In contrast, mice with high frac_up_, in which SST activation predominantly activated excitatory units (functionally IndNet-like), showed shorter intrinsic timescales during high than during low arousal.

**Fig. 6.**
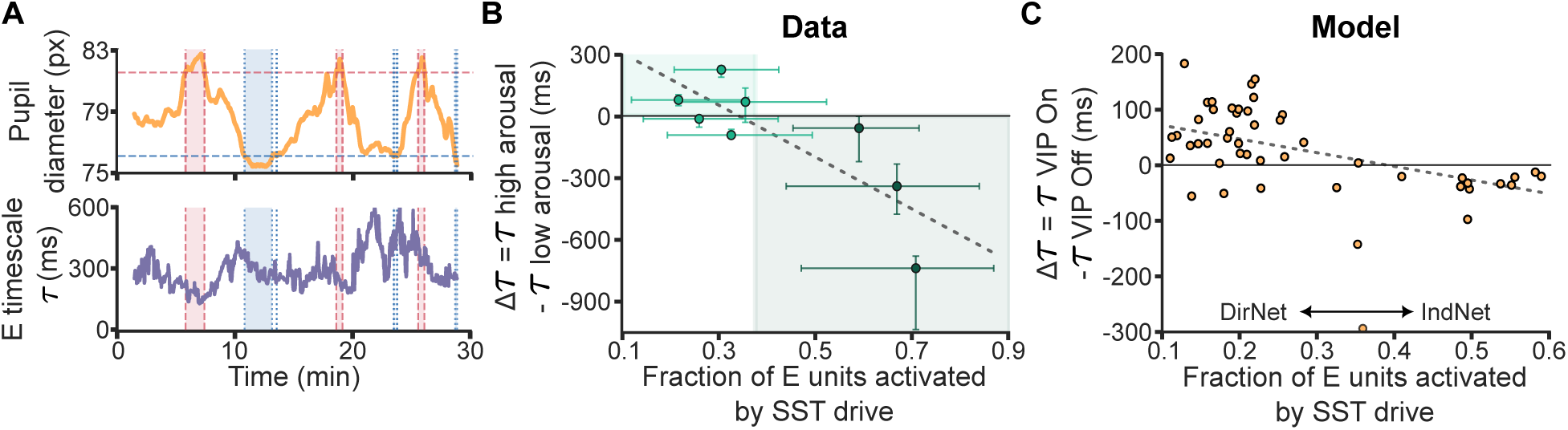
A negative correlation between the fraction of SST-activated excitatory neurons and arousal-dependent timescale changes. **A.** (Top) Pupil diameter (px, orange) for one example mouse during spontaneous activity. Horizontal dashed lines mark session-specific thresholds for high (red) and low (blue) pupil diameter. Vertical dashed lines and shaded bands mark the detected high-(red) and low-arousal (blue) bouts (see Methods). (Bottom) Median intrinsic timescale *τ* across V1 excitatory (E) units, computed in the same sliding windows, for the same mouse. **B.** Excitatory timescale difference between high and low arousal, Δ*τ* = *τ*_high_ *− τ*_low_, as a function of frac_up_, the fraction of E units activated by SST stimulation (*n* = 8 mice). Vertical error bars indicate bootstrap 95% confidence intervals (CIs) on Δ*τ*, and horizontal error bars indicate Wilson 95% CIs on frac_up_ (see Methods). The dashed line is a linear fit across mice (Pearson *r* = *−*0.811, *p* = 0.015; Spearman *ρ* = *−*0.786, *p* = 0.021). Shaded regions indicate the model-predicted ranges of timescale lengthening (light green, functionally DirNet-like) and shortening (dark green, functionally IndNet-like), separated at the model-derived zero crossing in **C**. **C.** Change in excitatory intrinsic timescale under VIP drive, Δ*τ* = *τ*_ON_ *− τ*_OFF_, as a function of frac_up_, measured separately during SST drive, for *n* = 50 networks spanning the IndNet-to-DirNet continuum (see Methods). The dashed line is a linear fit (Pearson *r* = *−*0.384, *p* = 0.006; Spearman *ρ* = *−*0.417, *p* = 0.003). Arrows indicate the directions toward the DirNet and IndNet endpoints.

We next asked whether the mixed-network model reproduced this empirical relationship. We treated VIP drive as an arousal-like modulation because arousal is known to preferentially recruit VIP INs across cortical areas [3, 15, 29, 61, 62]. For each network spanning the IndNet-to-DirNet continuum, we computed frac_up_ during SST drive and the change in excitatory intrinsic timescale, Δ*τ*, between conditions with and without VIP drive. Across networks, Δ*τ* decreased with frac_up_ (Fig. 6C; Pearson *r* = *−*0.384, *p* = 0.006; Spearman *ρ* = *−*0.417, *p* = 0.003). DirNet-like networks with low frac_up_ lengthened their intrinsic timescales under VIP drive, while IndNet-like networks with high frac_up_ shortened them.

In sum, these results suggest that the excitatory response to SST activation functionally predicts the direction of arousal-dependent timescale modulation across mice. The model predicts that varying SST-to-excitatory strength changes the relative influence of the direct and indirect SST pathways, which in turn leads to heterogeneous modulations of neuronal timescales and firing rates. The model thus identifies variation in SST-to-excitatory synaptic strength as a possible structural mechanism underlying the different functional response regimes observed across mice.

### Flexible arousal-like modulation of new assembly formation

We hypothesized that, depending on the relative influence of the two SST pathways, arousal-like VIP drive could either accelerate or slow the reorganization of existing assembly structure through synaptic plasticity. To test this hypothesis, we exposed networks with previously established excitatory assemblies to a new set of correlated inputs. We applied inputs targeting neuronal subgroups that differed from the original assemblies while synaptic plasticity remained active (Fig. 7A). After training, the network assembly structure rewired from the pre-trained assemblies to the new assemblies, the latter reflecting the new input correlations [50]. Examining the learning trajectories allowed us to measure how rapidly connections within the new subgroups strengthened and connections supporting the original assemblies weakened (Fig. 7B-C). In the presence of VIP drive, the two DirNet and IndNet network configurations exhibited opposite learning dynamics: VIP drive accelerated both the formation of new assemblies and the weakening of the original assemblies in DirNet (Fig. 7B), but slowed both processes in IndNet (Fig. 7C).

**Fig. 7.**
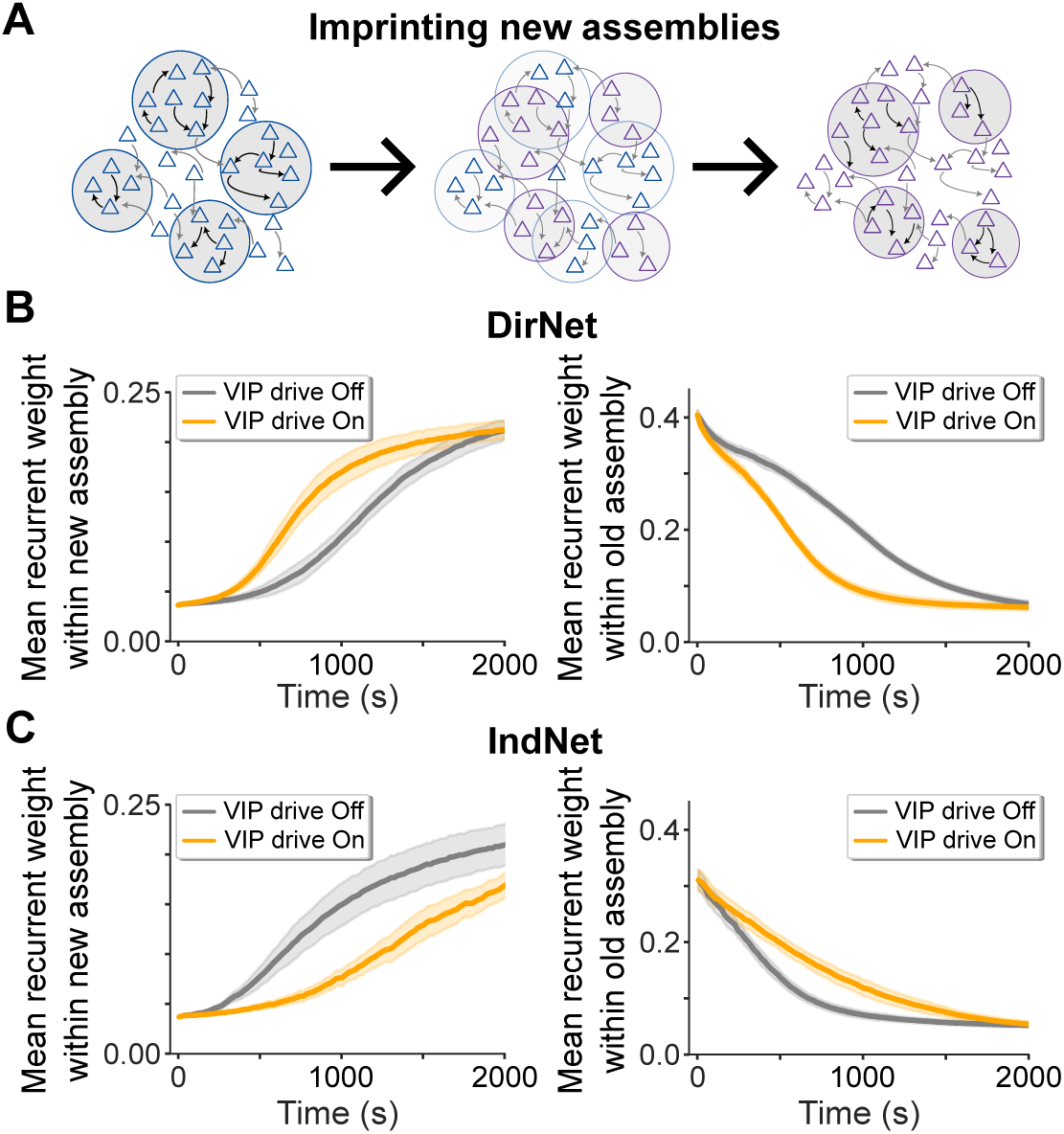
VIP drive accelerates or slows assembly reorganization. **A.** Schematic of the reorganization of the original assembly structure (blue) into a new assembly structure (purple) generated by repartitioning the excitatory neurons into new subgroups. **B.** (Left) Evolution of the mean excitatory-to-excitatory synaptic weight within the new assemblies in DirNet with (orange) and without (gray) VIP drive. (Right) Evolution of the mean excitatory-to-excitatory synaptic weight within the original assemblies under the same conditions. **C.** Same as **B** for IndNet.

Applying SST drive produced the mirror image of the effects of VIP drive on assembly formation (Fig. S5A–D). In DirNet, SST drive slowed the formation of new assemblies, while in IndNet it accelerated assembly formation. In the mixed network, the rate of assembly formation could likewise be shifted in either direction by selecting which SST subgroup was targeted (Fig. S5D). Across these manipulations, longer intrinsic timescales were associated with faster formation of new assembly structure, while shorter intrinsic timescales were associated with slower assembly formation.

Our results show that arousal-like VIP drive can flexibly modulate the formation of new assembly structure depending on the relative influence of the direct and indirect SST pathways. In DirNet, VIP drive accelerates assembly formation, whereas in IndNet it slows assembly formation. Together with the observed relationship between SST responses and arousal-dependent timescale changes across mice (Fig. 6B), these model results predict that mice with opposite timescale responses to arousal should also differ in how arousal affects the formation of new assemblies and the weakening of existing assemblies.

## Discussion

Although behavioral state strongly influences the temporal dynamics of cortical activity, the circuit mechanisms that account for the diversity of these effects across individuals have remained poorly understood. Analyzing publicly available *in vivo* recordings and optogenetic perturbations from the Allen Institute ‘Visual Coding’ dataset, we showed that the fraction of excitatory neurons activated by SST stimulation varied across mice and predicted whether intrinsic timescales increased or decreased during arousal. We hypothesized that this relationship could arise from differences in the relative influence of two pathways through which SST INs regulate excitatory activity: direct inhibition of excitatory neurons and indirect disinhibition through PV INs. By training networks with cell-type-specific synaptic plasticity, we generated circuits that exhibited similar spontaneous assembly dynamics despite differences in their SST-to-excitatory connectivity. In the model, arousal-like VIP drive lengthened intrinsic timescales and accelerated assembly reorganization when the direct pathway dominated and had the opposite effects when the indirect pathway dominated. Thus, even in circuits exhibiting similar spontaneous activity, the same modulatory signal can produce qualitatively opposite dynamical and computational consequences.

Previous research has mapped the landscape of intrinsic timescales across brain regions, characterized their diversity across individuals [6, 12], and analyzed these dynamics within the Allen Institute datasets [11, 21, 30]. In particular, locomotion has been found to modulate visual cortical dynamics during visual stimulation [21, 30]. Our analysis complements this work by relating timescale changes during pupil-defined arousal in the absence of visual stimulation to independently measured excitatory responses to SST stimulation. Our results thus suggest that cell-type-specific perturbations could provide a means of probing the circuit mechanisms underlying state-dependent cortical dynamics.

The variability in excitatory responses to SST stimulation across mice is consistent with differences in the functional organization of their local inhibitory circuits. In the model, varying the proportion of SST INs whose connections onto excitatory neurons strengthened during training was sufficient to reproduce the continuum of excitatory responses to SST activation and the negative relationship between the fraction of SST-activated excitatory neurons and arousal-dependent changes in intrinsic timescales observed across mice. Although the experimental measure distinguishes these different functional response regimes, identifying the anatomical or synaptic features that govern the position of an animal along this continuum will require further investigation. Synaptic strengths are often broadly distributed within cortical circuits [63], making substantial variability in the influence of SST-to-excitatory connections biologically plausible. Future work combining cell-type-specific physiology with connectomic measurements across animals could determine whether variation in SST-to-excitatory connectivity contributes to the functional response continuum observed here. Causal manipulations that shift the relative influence of the SST-to-excitatory and SST-to-PV-to-excitatory pathways should also shift both the excitatory response to SST activation and the timescale change produced by arousal-related input, providing a direct test of the mechanism proposed by the model.

Previous work established that top-down inputs targeting either excitatory or inhibitory neurons can alter the temporal dynamics of metastable activity in circuit models with excitatory and inhibitory populations [20, 30]. More recent studies incorporating multiple interneuron classes have shown that the same modulatory input can produce different effects depending on cell-type-specific connectivity, network state, and synaptic dynamics [31, 32]. These and other studies have highlighted the importance of direct inhibitory and indirect disinhibitory pathways in shaping circuit responses [33–35]. Our work extends these studies by showing how the relative influence of these pathways, established through cell-type-specific synaptic plasticity, determines the direction of modulation of metastable dynamics and intrinsic timescales. We focused on SST-to-excitatory connectivity because these synapses are known to undergo plastic changes [54]. However, differences in other synaptic connections could also shift the balance between direct inhibition and indirect disinhibition and generate similar functional regimes *in vivo*.

The opposite changes in intrinsic timescales were accompanied by differences in the rate of assembly reorganization. In the presence of arousal-like VIP drive, new assemblies formed more rapidly when direct SST-to-excitatory inhibition dominated, and more slowly when indirect disinhibition through PV INs dominated. These results suggest that differences in the relative influence of direct and indirect SST pathways can alter how rapidly a circuit’s existing assembly structure is modified. Because the formation of new assemblies relied on activity-dependent Hebbian plasticity, the changes in excitatory activity produced by VIP modulation likely contributed to the differences in learning dynamics. Thus, longer intrinsic timescales and faster assembly formation should be viewed as related consequences of the same circuit modulation rather than as evidence that the timescale change directly causes faster assembly formation. Together with the relation-ship between SST responses and arousal-dependent timescale changes observed *in vivo*, these model results predict that mice with opposite timescale responses to arousal should also differ in how arousal affects the reorganization of learned assembly structure.

Here, we focused on SST activation and arousal-related inputs to VIP. However, both interneuron classes are targets of diverse modulatory and long-range inputs. Their recruitment contributes to several computations, including locomotion-induced gain modulation [13–15], selective attention [23], context-dependent modulation [64, 65], and predictive processing [66]. These interneurons also contribute to novelty detection [67], flexible routing of information flow [68], regulation of global coherence [69], and gating of synaptic plasticity [70]. Other interneuron classes are also targets of modulatory inputs from motor and higher cortical areas. For instance, neuron-derived neurotrophic factor (NDNF)-expressing INs are located predominantly in layer 1 of the neocortex and inhibit several cortical cell types while largely sparing SST interneurons [71–73]. Pathways involving NDNF or other interneuron classes may further shape the effects of modulatory inputs. Nonetheless, our results show that differences in the relative influence of two pathways with opposing effects on excitatory activity are sufficient to produce opposing dynamical responses. The same framework could therefore be extended to other modulatory signals and circuit motifs in which direct inhibition competes with indirect disinhibition.

Several aspects of the experimental analysis should be considered when interpreting the model-data relationship. The proportions of excitatory neurons activated and suppressed by SST stimulation varied across cortical layers (Fig. S1A), and differences in laminar sampling across probes could therefore contribute to differences in frac_up_ across mice. Moreover, pupil-linked arousal covaries with locomotion and is associated with neuromodulatory signals that can recruit multiple cortical cell types. Changes in arousal may therefore engage multiple modulatory pathways, whereas the model delivers arousal-like input selectively to VIP INs. Although our analysis focused on spontaneous periods without visual stimulation, we could not isolate the contribution of any individual top-down or neuromodulatory pathway. Selective manipulation of SST or VIP INs during controlled changes in arousal would provide a more direct test of how these pathways shape intrinsic timescales and whether their effects depend on the functional response regime identified here.

Taken together, these results show that the same modulatory signal can drive cortical circuits toward opposite dynamical regimes depending on the relative influence of direct inhibition and indirect disinhibition. The excitatory response to SST activation may therefore provide a functional indication of the relative influence of these pathways and predict the direction of arousal-dependent changes in intrinsic timescales in individual animals.

## Methods

### Neuron and network model

We modeled a recurrent spiking network of the mouse cortical microcircuit, comprising four populations of neurons: one excitatory and three inhibitory interneuron populations (Fig. 2A). The network consisted of a total of 4000 excitatory neurons and 1000 inhibitory interneurons (500 PV, 250 SST and 250 VIP). Single neurons were modeled as current-based leaky integrate-and-fire neurons with the membrane voltage dynamics:

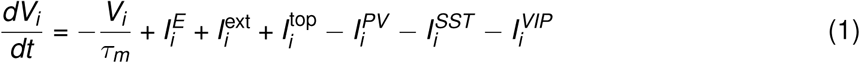

where *τ_m_* is the membrane time constant. The input current for each neuron was defined as the signed sum over all input sources, which included external feedforward and top-down (when present) excitatory inputs and recurrent synaptic inputs from other neurons in the network (excitatory, PV, SST and VIP). All weights *w_ij_* are defined with positive magnitudes (Table T1). Upon reaching the spiking threshold *V_thr_*, the neuron elicited a spike and the membrane potential was reset to *V_res_*, where it was clamped for an absolute refractory period *τ_abs_*. Connections between neurons of different populations were constrained by experimental findings [25, 74]. Excitatory neurons targeted all other neuron populations and were inhibited by PV and SST INs. VIP INs inhibited SST INs which in turn inhibited PV and VIP INs. PV INs targeted other PV and VIP INs. For the interacting neuron types, connections were chosen randomly with probability 0.1, with autapses being removed afterwards (Table T1). Each synaptic current decayed exponentially between spikes and increased by *w_ij_*/τ*^X^*_syn_ at each spike of a presynaptic neuron *j* in population *X* :

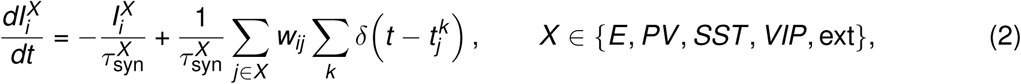

where *t_j_^k^* is the *k* -th spike time of neuron *j*, and 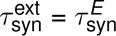.

### Excitatory plasticity

Synapses between excitatory neurons were plastic according to a triplet STDP rule [49]. Each excitatory neuron *j* carried two presynaptic traces, *r*_1,*j*_ and *r*_2,*j*_, and two postsynaptic traces, *o*_1,*j*_ and *o*_2,*j*_, which decayed exponentially and increased by 1 at each of its spikes:

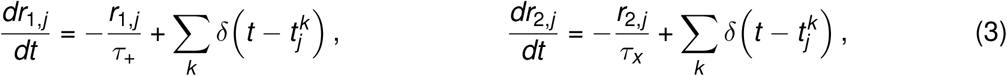

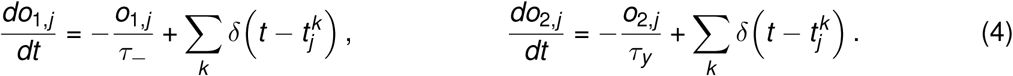

The weight *w_ij_* of the synapse from *j* to *i* changed only at spike times of *j* or *i*:

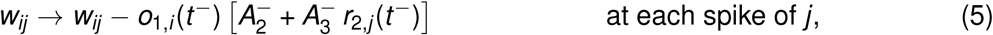

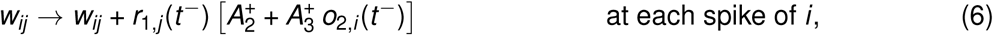

and was then clipped to 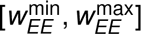 (Table T1). Here *t^−^* denotes the value immediately before the spike: at each spike, the weights were updated first and the traces of the spiking neuron were incremented afterwards, so a spike did not contribute to its own triplet term. The amplitudes A_2_^+^, A_2_^−^, A_2_^+^, A_2_^−^ (Table T2) determined the amount of synaptic potentiation (+) or depression (*−*) resulting from pairs (‘2^′^) and triplets (‘3^′^) of spikes. Pre-post pairs and (1-pre)-(2-post) triplets induced potentiation, post-pre pairs and (2-pre)-(1-post) triplets led to depression of the respective synapses. To avoid saturating the synaptic weights, we included heterosynaptic competition between excitatory synapses that targeted the same postsynaptic excitatory neuron [50, 75]. Every Δ*t*_norm_ = 20 ms, the incoming E-to-E weights of each neuron *i* were shifted,

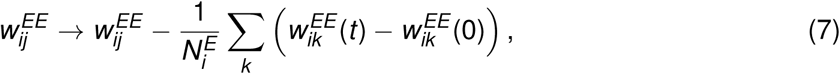

with the sum over the *N_i_^E^* existing synapses onto *i*, and then clipped to 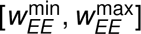. Clipping can break exact conservation, so the summed weight is approximately constant.

### PV inhibitory plasticity

PV-to-excitatory inhibitory synapses were governed by a pair-based inhibitory STDP rule (iSTDP) which homeostatically regulates excitatory firing [52]. Analogous to the triplet STDP, each PV neuron *j* carried a presynaptic trace 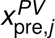 and each excitatory neuron *i* a postsynaptic trace 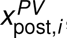, both with time constant *τ_PV_* :

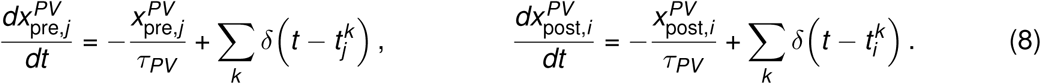

The PV-to-E weights were updated as

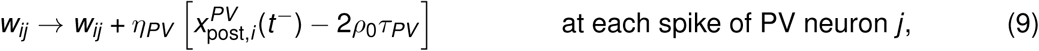

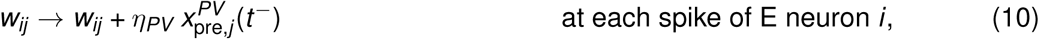

and clipped to 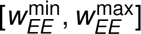, with the same update order as for E-to-E plasticity. The parameter *η_PV_*, called learning rate, scaled the overall amount of long-term plasticity elicited by spike pairs, similar to the amplitudes in the triplet STDP. The parameter *ρ*_0_ denoted a target set-point for the firing rate of the postsynaptic excitatory neuron. A presynaptic spike that was not paired with a postsynaptic spike led to the depression of the synaptic weight by 2*ρ*_0_*τ_PV_ η_PV_* . When a presynaptic spike and a postsynaptic spike occured in close temporal proximity, the extent of potentiation was determined by the time difference between the pre- and postsynaptic spike.

### SST inhibitory plasticity

SST-to-excitatory inhibitory synapses were governed by an asymmetric spike-timing-dependent plasticity rule (STDP) [54, 76]. Unlike symmetric STDP models, this rule specifically amplified the causal relationship between neuronal firing, using distinct amplitudes for potentiation and depression. The dynamics of the synaptic traces were represented by decaying exponentials updated upon spike occurrences:

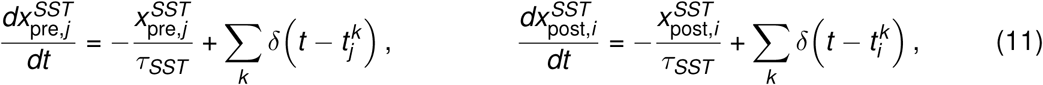

for each SST neuron *j* and excitatory neuron *i*. The SST-to-E weights were updated as

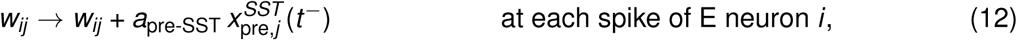

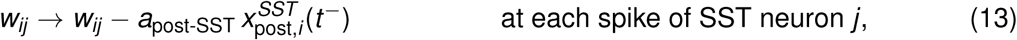

and clipped to 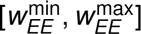, with the same update order as above. Here, *a_pre_*_−*SST*_ and *a_post_*_−*SST*_ denoted the learning rates for potentiation and depression, respectively, establishing the asymmetric nature of synaptic adjustments. Potentiation predominantly occurred when a presynaptic spike preceded a postsynaptic response within a critical time window, while depression was more likely when the postsynaptic spike preceded the presynaptic event, reinforcing the directionality in synaptic modifications.

### Stimulation protocol for assembly formation

To induce the formation of neuronal assemblies, excitatory neurons were partitioned into an unclustered background (1000 neurons) and a clustered (3000 neurons) group. Within the clustered group, neurons were further divided into 15 subgroups containing 200 neurons each. All neurons received a constant feedforward input in the form of spike trains that followed Poisson statistics with a rate of 5kHz. After 10s of thermalization, all 15 subgroups consecutively and randomly received additional feedforward input for 1s, followed by 1s break. The stimulation protocol above was repeated for *N_trials_* = 18 (540*s*). The simulation continued with plasticity on and no patterned input until 800s (Fig. S3, S4). All later simulations started from the 570s snapshot. In the same way, the synaptic weights and dynamical state variables used in the top-down modulation sections were taken at *t* = 570s, i.e. 20s after the last stimulus. Each network instance was initialized with an independent random connectivity, initial condition and stimulation order.

### Top-down modulation

For studying top-down modulation, we inactivated both excitatory and inhibitory plasticity 20s after training (at 570s), given that modulatory effects act over shorter timescales than those governed by plasticity [77]. Inactivation of iSTDP was necessary because the iSTDP rule prevents lasting changes in excitatory firing rates [50, 52] which we wanted to include to study the effects of top-down modulation. Inactivation of the plasticity between excitatory neurons was necessary because the network with excitatory plasticity is unstable without firing homeostasis [50, 78]. Each modulation simulation was restarted from the network state saved at 570 s (synaptic weights, membrane potentials, synaptic currents and plasticity traces). At modulation onset (*t* = 1 s), each neuron *i* of the targeted population (VIP or SST) was assigned a constant offset current *I*^top^, drawn once and independently for each neuron from a normal distribution *N* (0, *σ*^2^) and held fixed for the rest of the simulation; neurons of non-targeted populations had *li^top^* = 0. The offset enters the membrane equation directly as an additional input current (Eq. 1), without synaptic filtering. The standard deviation was *σ* = *r_X_ I*_baseline,*X*_, where *I*_baseline,*X*_ = *w_XF_ r*_ext_ is the mean feedforward input current to population *X* and *r_X_*is the modulation strength (*r_S_* for SST, *r_V_*for VIP; Table T1). Because the offsets have zero mean, the modulation leaves the population-averaged input unchanged and increases the heterogeneity of inputs across neurons; a larger *r_X_* corresponds to stronger top-down input [20, 30]. Unmodulated control simulations (*r_X_* = 0) were started from the same saved state. Qualitatively similar results can be obtained with a constant positive offset current applied to all targeted neurons.

### Analysis of metastable activity

To characterize metastable activity in the spiking network, we computed, for each assembly, the population firing rate of its 200 neurons in a 20ms sliding window advanced in steps of 0.1ms. At each time step we retained the rate of the most active assembly and its identity, and smoothed this maximum rate with an additional 10 ms moving average. Periods during which the smoothed rate exceeded *f*_thr_ = 10 Hz were defined as assembly activations and assigned to the assembly that was most often the most active during that period; consecutive activations assigned to the same assembly and separated by less than 100 ms were merged. Active times were the durations of the activations assigned to each assembly. Inactive times were the intervals between two consecutive activations of the same assembly. We did not observe intervals during in which multiple assemblies were active at the same time. Durations shorter than the sliding window value were excluded from the distributions in Figs. 3C,D and 4C,D.

### Mean-field approximation

Mean-field theory was used in a simplified network of *N* = 800 neurons with two excitatory assemblies (Table T3 with synaptic weights scaled by Scale = 5000*/N* = 2.5) using the population density approach [20, 79]. This is a separately parameterized reduced model chosen to reproduce the DirNet and IndNet regimes. Synapses within an assembly were potentiated by a factor *J*_+_ = 3.1 and synapses between assemblies depressed by *J*_−_ = 1 *− γf* (*J*_+_ *−* 1) with *γ* = 0.5 [80]. Firing thresholds *θ_α_*were determined numerically for each network such that the spontaneous rates *r_E_* and *r_γ_* of Table T3 formed a stable fixed point of the unclustered network (*J*_+_ = 1). The input to each neuron was completely characterized by the infinitesimal mean *µ_α_* and variance *σ*^2^ of the postsynaptic current for *Q* + 4 neural populations: the first *Q* populations representing the *Q* excitatory clusters, the (*Q* + 1)th population representing the background unclustered excitatory population, and the remaining populations representing the inhibitory population (PV, SST and VIP, respectively).

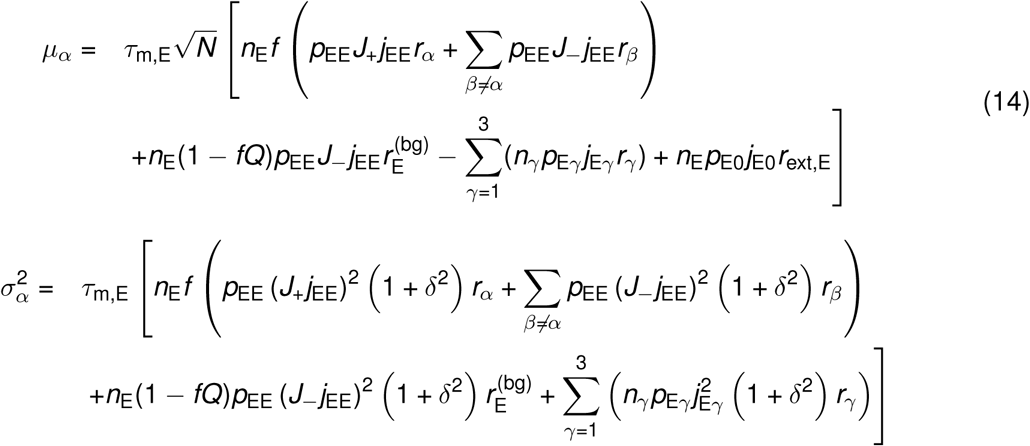

where *r_α_*, *r_β_*, with *α*, *β* = 1, …, *Q*, were the E-cluster firing rates; *r_E_*^(bg)^ was the background E-population firing rate; *r_γ_* was the *γ*-population firing rate with *γ ∈ {PV*, *SST*, *VIP}*; *n*_E_, *n_γ_* were the fractions of E and I neurons, respectively; and *δ* = 0 was the standard deviation of the synaptic weight distribution. The two sources contributing to the variance *σ*^2^ were thus the temporal variability in the input spike trains from the presynaptic neurons, and the quenched variability in the synaptic weights and connectivity. The afferent current to a background E-population neuron read:

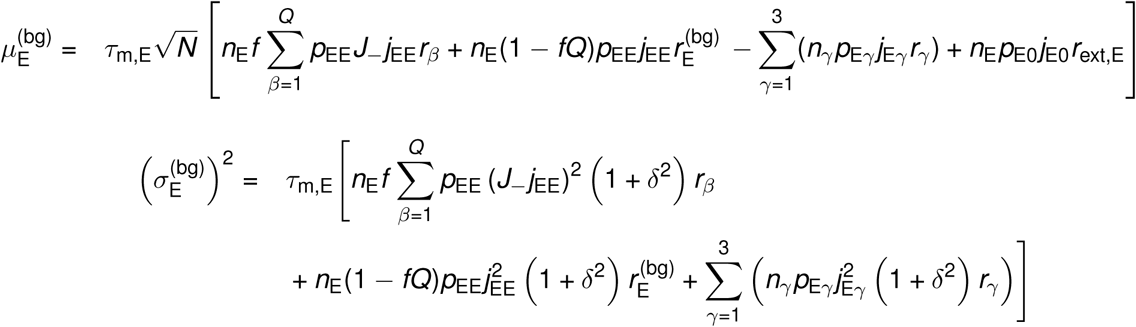

and, similarly, for each I population *γ*:

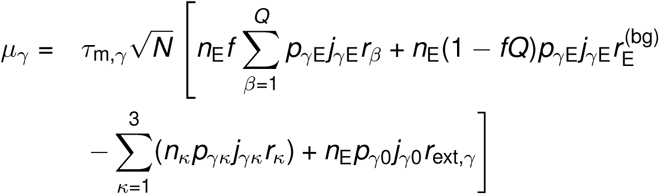

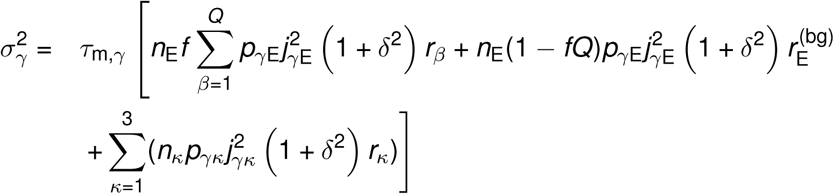

The network fixed points satisfied the *Q* + 4 self-consistent mean-field equations [80]:

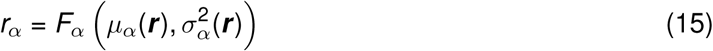

where 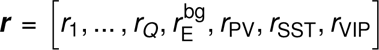 was the population firing rate vector (boldface represents vectors). *F_α_*was the current-to-rate function for population *α*, which varied depending on the population and the condition. In the absence of the top-down modulation, the LIF current-to-rate function *F* ^0^ was used:

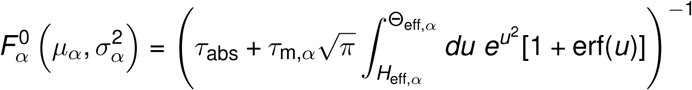

where Θ_eff,*α*_ =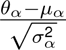 + *ak_α_*, H_eff,*α*_ = 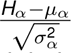 + *ak_α_*. Here, *k_α_* = 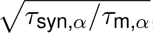, *a* = 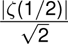 *≈* 1.03 [81, 82]. In the presence of top-down modulation, the current-to-rate function of the targeted population (in our case, *α* = *Q* +3 or *α* = *Q* +4) was evaluated at a shifted mean afferent current, 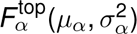 = 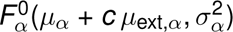, where *µ*_ext,*α*_ = *I*_0_ *≡ τ_m_*_,*α*_√*N n_E_ p_α_*_0_ *j_α_*_0_ *r*_ext,*α*_ was the baseline (unmodulated) afferent current already appearing in *µ_α_* (Eq. 14) and *c* was the top-down modulation strength, expressed as a fractional increase of *µ*_ext,*α*_, playing an analogous role of *r_S_*and *r_V_* in the spiking model. Top-down modulation thus acted as a deterministic increase in the mean input current to the targeted population; it did not contribute additional input variance *S*_α_^2^. Qualitatively similar results can be obtained with an additional input variance. Fixed points *r*\* of Eq. 15 were found with Newton’s method; the fixed points were stable (attractors) when the reduced stability matrix with input variance held fixed

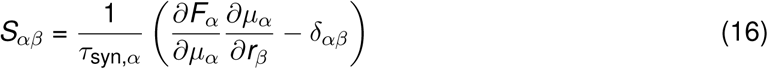

evaluated at *r*\* had only eigenvalues with negative real parts. Stability was defined with respect to an approximate linearized dynamics of the mean *m_α_*and variance *S*_α_^2^ of the input currents [83]

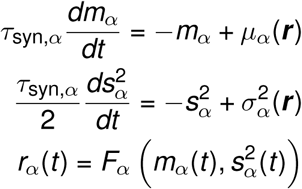

where *µ_α_* and *σ*^2^ were the stationary values given above [20].

The mean-field equation (Eq. 15) for the *P* = *Q* + 4 populations may be reduced to a set of effective equations governing the dynamics of a smaller subset of *q < P* populations, henceforth referred to as populations in focus [84]. The reduction was achieved by integrating out the remaining *P − q* out-of-focus populations. This procedure was used to estimate the energy barrier separating the two network attractors in Fig. 3 and Fig. 4. Given a fixed set of values ***~r*** = [*~r*_1_, …, *~r_q_*] for the in-focus populations, one solved the mean-field equations for *P − q* out-of-focus populations

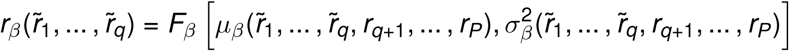

for *β* = *q* + 1, …, *P* to obtain the stable fixed point ***r*** *^f^* (***~r***) =. [*r_q_*_+1_(***~r***), …, *r_P_*(***~r***)] of the out-of-focus populations as functions of the in-focus firing rates ***~r***. Stability of the solution ***r*** *^f^* (***~r***) was computed with respect to the stability matrix of the reduced system of *P −q* out-of-focus populations. Substituting the values ***r*** *^f^* (***~r***) into the fixed-point equations for the *q* populations in focus yielded a new set of equations relating input rates ***~r*** to output rates ***r*** ^out^:

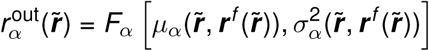

for *α* = 1, …, *q*. The input ***~r*** and output ***r*** ^out^ firing rates of the in-focus populations will be different, except at a fixed point of the full system where they coincide. The correspondence between input and output rates of in-focus populations defined the effective current-to-rate transfer functions

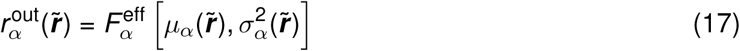

for *α* = 1, …, *q* in-focus populations at the point ∼*r*. The fixed points 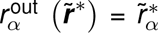 of the in-focus equation (Eq. 17) were fixed points of the entire system. It may occur, in general, that the out-of-focus populations attained multiple attractors for a given value of ∼*r*, in which case the set of effective transfer functions *Fα*^eff^ was labeled by the chosen attractor; in our analysis of the two-clustered network, only one attractor was present for a given value of ***~r***.

### Energy potential

In a network with *Q* = 2 clusters, one could integrate out all populations (out-of-focus) except one (in-focus) to obtain the effective transfer functions for the in-focus population representing a single cluster, with firing rate ∼*r* (Eq. 17 for *q* = 1). Network dynamics can be visualized on a one-dimensional curve, where it was well approximated by the first-order dynamics (see [84] for details):

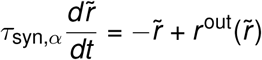

These dynamics can be expressed in terms of an effective energy function *E* (*~r*) as

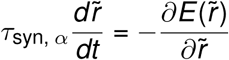

so that the dynamics can be understood as a motion in an effective potential energy landscape, as if driven by an effective force 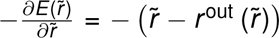. The minima of the energy with respect to *~r* were the stable fixed points of the effective one-dimensional dynamics, whereas its maxima represented the effective energy barriers between two minima. The one-dimensional reduction of the two-cluster network had three fixed points: a low-rate stable state (assembly inactive), a high-rate stable state (assembly active), and an unstable fixed point *C* between them. We computed Δ (Eq. 18) with *A* the high-rate state, so Δ sets the escape rate from the active state:

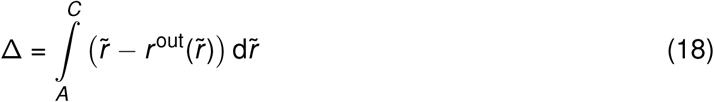

which represented the area between the identity line (*y* = *~r*) and the effective transfer function (*y* = *r* ^out^(*~r*)). In the finite network, where the dynamics comprised stochastic transitions among the states, switching between the two stable points would occur with a frequency that depended on the effective energy barrier.

### Combined mixed networks

We split the SST population into two subgroups during training: the former projecting plastic connections and the latter static, weak connections to the excitatory neurons (Fig. 5A). The training protocol mirrored the one employed in DirNet and IndNet networks. We co-activated non-overlapping subgroups of neurons in random order during training to form neuronal assemblies via synaptic plasticity (Fig. 5A). Despite the partitioning of SST INs into two subgroups, the weight evolution within individual E neuron subgroups was congruent with the previous networks (Fig. S4, compared to Fig. S3). After training, each SST subgroup received top-down inputs drawn from a normal distribution similar to the DirNet and IndNet case.

### Allen Institute Neuropixels dataset and unit selection

We analyzed the Allen Institute ‘Visual Coding’ Neuropixels dataset [37], in which each recording session corresponds to a different mouse, with six cortical visual areas (V1/VISp, LM/VISl, AL/VISal, RL/VISrl, AM/VISam, PM/VISpm) and the thalamic nuclei LGN and LP recorded simultaneously (Fig. 1A). Spike times were extracted with the Allen Software Development Kit (AllenSDK). All quantitative analyses below (unit classification, arousal bouts, intrinsic timescales) were restricted to the primary visual area (V1/VISp). For cell-type-specific optogenetic stimulation, the Allen Institute crossed Sst-IRES-Cre and Pvalb-IRES-Cre driver lines with the Ai32 reporter line to express channelrhodopsin-2 (ChR2) in SST and PV INs, respectively. We refer to these mice as Sst-Cre and Pvalb-Cre mice throughout.

To identify optotagged interneurons, we used the 10 ms square-pulse condition (selected as epochs of 9–20 ms duration in the AllenSDK optogenetic stimulation table) delivered to *Sst* -Cre and, for a specificity control (see below), *Pvalb*-Cre mice, and computed, for every unit recorded in VISp (spike counts binned at 0.5ms), the mean firing rate in an 8ms baseline window (*−*10 to *−*2ms relative to pulse onset) and an 8ms evoked window (+1 to +9ms relative to pulse onset).

Trials at all light levels were pooled. A unit was classified as optotagged if

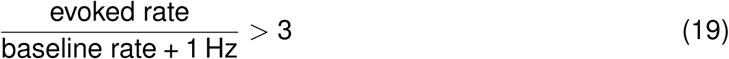

where the additive 1Hz term regularizes the ratio for low-firing-rate units. Of the 58 mice in the dataset, 12 contained *Sst* -Cre optotagged units recorded simultaneously in VISp and 8 contained *Pvalb*-Cre optotagged units recorded simultaneously in VISp.

Excitatory (E) units were defined, within VISp, as units with amplitude cutoff *≤* 0.01, waveform duration *≥* 0.35ms (broad-spiking, used as a proxy for putative excitatory/regular-spiking cells), and excluding optotagged units, yielding 392 E units across the 12 *Sst* -Cre mice and 179 E units across the 8 *Pvalb*-Cre mice. Of the 12 *Sst* -Cre mice, 11 had at least 6 E units, 10 of these had eye-tracking data, 9 yielded a computable timescale difference, and 8 passed the locomotion criterion below. All per-mouse results (Fig. 1E, Fig. S1B,C and Fig. 6B) are shown for these 8 mice, so that each mouse’s frac_up_ in Fig. 1E is the value on the horizontal axis of Fig. 6B. Of the 8 *Pvalb*-Cre mice, 6 had eye-tracking data and 5 entered the specificity control below (Fig. S1D, Fig. S6). To rule out a contribution of light reaching the retina during optogenetic stimulation, we applied the same before/during comparison to the full-field visual flash trials (*∼* 250 ms, all colors pooled) of each mouse, giving a flash-evoked Δrate for every E unit. Across E units of the *Sst* -Cre cohort, opto- and flash-evoked Δrate were only weakly correlated (Pearson *r* = 0.15, *n* = 392 units from 12 mice), indicating that the response to SST activation was not primarily driven by a light artifact. As a robustness check, we labeled a unit flash-like if its opto-evoked Δrate was within 2 Hz of its flash-evoked Δrate, removed all flash-like units, and recomputed frac_up_ and the population timescale from the remaining units; the relationship between frac_up_ and Δ*τ* was preserved (Pearson *r* = *−*0.77, *p* = 0.026; Spearman *ρ* = *−*0.76, *p* = 0.028; *n* = 8 mice). One *Pvalb*-Cre mouse was excluded from the PV specificity control since none of its 20 E units responded significantly to PV activation (paired Wilcoxon test, *p <* 0.05), and 19 of the 20 were flash-like by the criterion above.

For each mouse, spontaneous activity (low-luminance gray screen, no visual stimulation) was identified for the timescale analysis below. If the single longest continuous spontaneous block exceeded 400s (*≈* 6.7 min), it alone was used; otherwise, every spontaneous epoch of at least 120s was analyzed independently and its sliding windows pooled, so that no analysis window spanned a gap between epochs. Pooling across both cases, 11 of the 12 *Sst* -Cre mice and all 8 *Pvalb*-Cre mice contributed valid intrinsic-timescale estimates.

Mice that spent *≥* 98% of the recorded session moving (duration-weighted fraction of time with |running speed*| >* 2cm/s) were excluded from the arousal analysis, as too little quiescent time remained to estimate a low-arousal baseline (1 *Sst* -Cre mouse excluded; no *Pvalb*-Cre mice excluded).

Cortical layer assignment (Fig. S1) was performed at the level of recording channels: each channel’s CCF (Allen Common Coordinate Framework) coordinate was mapped to the corresponding structure annotation in the 10 *µ*m-resolution CCF annotation volume, and a layer identifier was extracted from the resulting structure acronym (e.g. ”VISp5” *→* layer 5, ”VISp2/3” *→* layer 2/3). Units were assigned the layer of their peak recording channel; channels without a resolvable cortical-layer annotation were excluded from the layer-resolved analysis.

### Excitatory-unit responses to optogenetic SST activation

To classify each V1 excitatory unit’s response to SST activation (Fig. 1D-E), we considered opto-genetic stimulation epochs of nominal duration 1 s (the ”raised cosine” and ”fast pulses” conditions; the brief 5/10 ms pulses used for optotagging above were too short for a reliable rate estimate), yielding on average *∼*90 trials per session. For each trial and each E unit, we computed the firing rate in an equal-duration window immediately before the stimulus ([start *−* duration, start)) and during the stimulus ([start, stop)), and defined Δrate = rate_during_ *−* rate_before_.

For each mouse, we defined

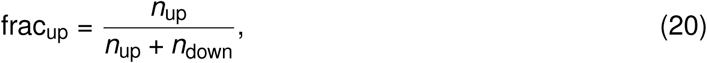

where *n*_up_ and *n*_down_ are the number of that mouse’s E units with Δrate *>* 0 (activated by SST drive) and Δrate *<* 0 (suppressed by SST drive), respectively (Fig. 1E, Fig. S1). Units with Δrate = 0 were counted as neither activated nor suppressed and excluded from the denominator. The same definition was used for all per-mouse fractions (Fig. 1E, Fig. S1C,D, Fig. 6B, Fig. S6) and for the model networks (Fig. 5D, Fig. 6C). Units suppressed by SST drive can be described with a strong, effective SST-to-excitatory connection (the DirNet configuration in our model), while units activated by SST drive can be described with a weak SST-to-excitatory connection in which PV-mediated disinhibition dominates (the IndNet configuration). Mice with low frac_up_ are therefore DirNet-like and mice with high frac_up_ are IndNet-like.

### Arousal bouts from pupil diameter

We measured pupil size from the Allen Institute’s eye-tracking output as the effective diameter 2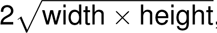, discarding samples flagged as blinks or otherwise missing. For each of the 120s sliding windows to evaluate intrinsic timescales (see below), pupil diameter was averaged over [window center *−* 60s, window center + 60s) (Fig. 6A, top).

Within each session, ”high” and ”low” arousal windows were defined as those whose pupil diameter exceeded that session’s own 90th percentile or fell below its own 10th percentile, respectively, with percentiles computed across all windows in that session. A mouse was included in the arousal analysis only if it had at least 5 qualifying high windows and 5 qualifying low windows.

### Intrinsic timescale analysis

For both the model and the data, intrinsic timescales were estimated from single-neuron spike-count autocorrelations computed in Δ*t* = 30ms bins. Within an analysis window of *N* bins, the spike-count sequence 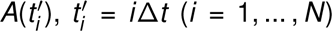, was z-scored using its window-wide sample mean 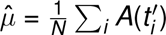 and standard deviation 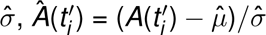, and the autocorrelation at lag *t_j_* = *j*Δ*t* was computed as

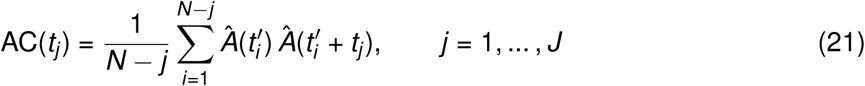

up to a maximum lag of 1s.

The autocorrelation was fit with an exponential decay function using nonlinear least squares

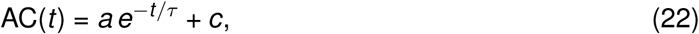

where *τ* is the estimated intrinsic timescale. Lag 0 was excluded. During fitting, parameters were constrained to 0.001 *≤ a ≤* 1.01, *−*0.1 *≤ c ≤* 0.1 and *τ ≥* 0.03 ms, with initial values *a*_0_ = max*_j_* AC(*t_j_*) *−* min*_j_* AC(*t_j_*), *τ*_0_ = 300 ms and *c*_0_ = min*_j_* AC(*t_j_*). Fits that failed to converge were discarded.

For the data, autocorrelations were computed in a sliding window of duration 120 s, advanced in steps of 2 s across the retained spontaneous blocks; the resulting fit was retained only if *R*^2^ *≥* 0.5 and *τ >* 0. The population timescale for a window was the median *τ* across all qualifying E units in that window (Fig. 6A, bottom). For each mouse, Δ*τ* = *τ*_high_ *− τ*_low_, where *τ*_high_ (*τ*_low_) was the median population timescale across that mouse’s high-(low-) arousal windows (Fig. 6B). A 95% confidence interval on Δ*τ* was obtained by bootstrap: the high- and low-arousal window sets were resampled independently with replacement 2000 times, the difference of medians recomputed on each resample, and the interval taken as the 2.5th/97.5th percentile of the resulting distribution. Consecutive windows overlap by 118 s and are therefore not independent. A bootstrap over arousal bouts (runs of consecutive qualifying windows) was not possible because 4 of the 8 mice had fewer than 3 independent bouts in at least one arousal condition. These bootstrap intervals therefore do not account for temporal dependence between windows and were not used in any statistical test. Across mice, Δ*τ* and frac_up_ were related by ordinary least-squares linear regression, and by Pearson and Spearman correlation (Fig. 6B); these used one estimate of Δ*τ* per mouse and did not treat windows as independent observations. The 95% interval on each mouse’s frac_up_ is the Wilson score interval for a binomial proportion.

For the model, autocorrelations were computed over each simulation’s entire modulation window (from the onset of top-down modulation to the end of the simulation), independently for the unmodulated and modulated conditions, using the same 30ms bins and exponential fit as above. A neuron’s fit was retained only if: (1) *R*^2^ *≥* 0.5; (2) the autocorrelation decreased monotonically over its first two lags; (3) its firing rate was *≥* 0.01Hz; and (4) it passed criteria (1)–(3) in *both* the unmodulated and modulated conditions (paired comparison). The network timescale for a given condition was defined as the median of the resulting *τ_i_* distribution across all surviving excitatory neurons.

### Direct-to-indirect ratio in the model

To relate frac_up_ to Δ*τ* in the model (Fig. 6C), we used five connectivity configurations spanning the IndNet-to-DirNet continuum – pdi = 0 (IndNet), 0.25, 0.5, 0.75, and 1 (DirNet), where pdi is the fraction of SST INs projecting strong (plastic) connections onto excitatory neurons (during training), each simulated in 10 independently trained networks (*n* = 50 networks total).

frac_up_ was computed as in the data (above). E-neuron firing rates were binned in 5s windows with and without SST drive, each neuron was labeled by the sign of its rate change, and frac_up_ was the fraction of neurons with a positive rate change. In contrast, Δ*τ* was computed under VIP drive, motivated by arousal preferentially recruiting VIP interneurons [62]. Therefore, Δ*τ* = *~τ*_ON_ *− ~τ*_OFF_, where *~τ* is the median timescale (computed as described above) across excitatory neurons with and without VIP drive.

Across the 50 networks, Δ*τ* and frac_up_ were related by ordinary least-squares linear regression, and by Pearson and Spearman correlation (Fig. 6C). The boundary frac_up_ = 0.388 shown in Fig. 6B is the zero-crossing of this linear fit for the model, marking the transition between DirNet-dominated (predicted Δ*τ >* 0) and IndNet-dominated (predicted Δ*τ <* 0) regimes.

### Learning speed analysis

To measure how quickly a novel assembly structure could be acquired under top-down modulation (Fig. 7A–C, Fig. S5), each trained network was resumed from its post-training synaptic weight configuration (saved at 570s, see above) together with all dynamical state variables (membrane potentials, synaptic currents, and plasticity traces). The 3000 excitatory neurons originally assigned to the 15 assemblies were reshuffled, independently for each network instance, into 15 new non-overlapping subgroups of 200 neurons each. The 1000 unclustered background neurons were not included in this re-partitioning. Because reassignment was random, each new subgroup shared on average 200 *×* 200*/*3000 *≈* 13 neurons with any given original assembly. Excitatory, PV-to-excitatory and SST-to-excitatory plasticity remained active throughout re-stimulation. SST-to-E plasticity was active only where it had been enabled during initial training. Following a 100ms thermalization period, the 15 new subgroups were stimulated consecutively in random order with the same additional feedforward Poisson drive used during initial training (see ”Stimulation protocol for assembly formation”), but with a shorter stimulus duration (200ms) and inter-stimulus gap (400ms). This protocol was repeated for *N_trials_* = 240. The modulatory input (SST or VIP drive) was added at the onset of each stimulus and withdrawn at its offset, in contrast to the tonic modulation held throughout the standard modulation experiments (see ”Top-down modulation”). Synaptic weight matrices were saved every 20s throughout re-stimulation. We tracked the evolution of the mean synaptic weight within the new (rearranged) assemblies and within the original assemblies over the course of re-stimulation, with and without top-down modulation.

### Network simulations

All model simulations were performed using custom software written in Julia and Python 3.11. Each network was initialized with random synaptic connections and simulated with random initial conditions. Dynamical equations for the leaky integrate-and-fire neurons were integrated with the forward Euler method with Δ*t* = 0.1ms. Mean-field analysis was performed using MATLAB 2025b.

## Data availability

The Allen Brain Observatory Visual Coding Neuropixels data are publicly available through the AllenSDK [37]. Simulation and analysis code is available at https://github.com/comp-neural-circuits/ top-down-timescales.

## Contributions

F.V. and J.G. designed research; F.V. and L.M. performed the mean-field analysis; F.V. performed the numerical simulations; F.V. analyzed the Neuropixels data; F.V., L.M. and J.G. wrote the paper.

## Acknowledgments

This work has received funding from the European Research Council (ERC) under the European Union’s Horizon Europe research and innovation programme (ERC CoG FeedbackCircuits, grant agreement no. 101170267). We thank Irina Pochinok and all members of the “Computation in Neural Circuits” group for helpful discussions.

## Supplementary Material

**Fig. S1.**
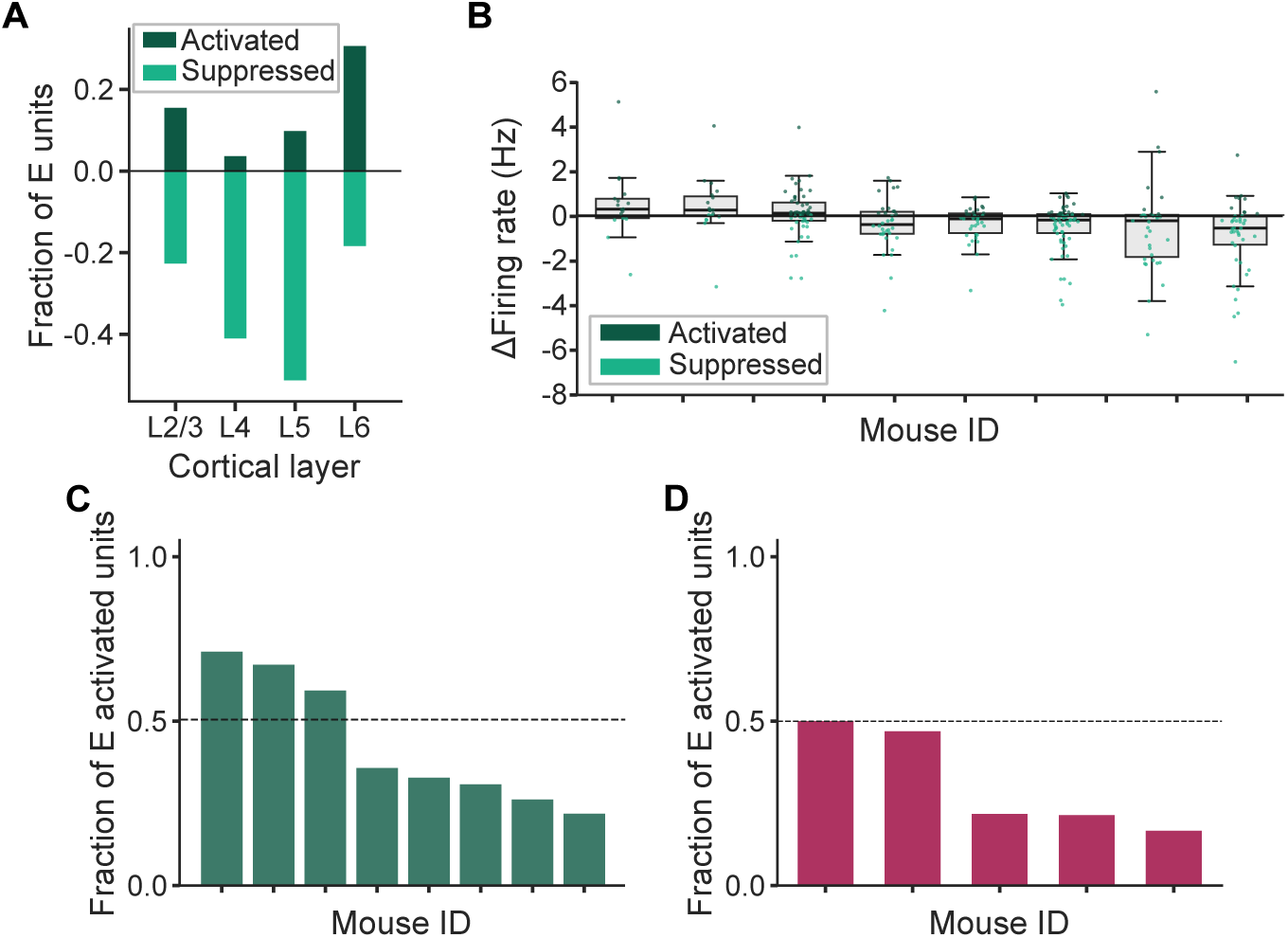
Layer dependence, per-mouse distributions and cell-type specificity of excitatory-unit responses to optogenetic interneuron activation. **A.** Fraction of layer-matched V1 excitatory (E) units activated (dark green) or suppressed (light green) by optogenetic SST drive, grouped by the cortical layer of the recorded interneuron (L2/3, L4, L5, L6; see Methods). **B.** Distribution of firing-rate changes (Δfiring rate, Hz) of individual V1 E units in response to optogenetic SST activation, for each *Sst* -Cre mouse, split by response sign (dark green: activated; light green: suppressed; see Methods). **C.** Fraction of E units activated by SST drive, frac_up_, for each *Sst* -Cre mouse (*n* = 8), computed from the unit counts in Fig. 1E and used as the horizontal axis of Fig. 6B. The dashed line marks frac_up_ = 0.5, above which activated units outnumber suppressed ones. **D.** Same as **C.** for E units activated by optogenetic PV drive in *Pvalb*-Cre mice (*n* = 5), the cohort used as a specificity control in Fig. S6.

**Fig. S2.**
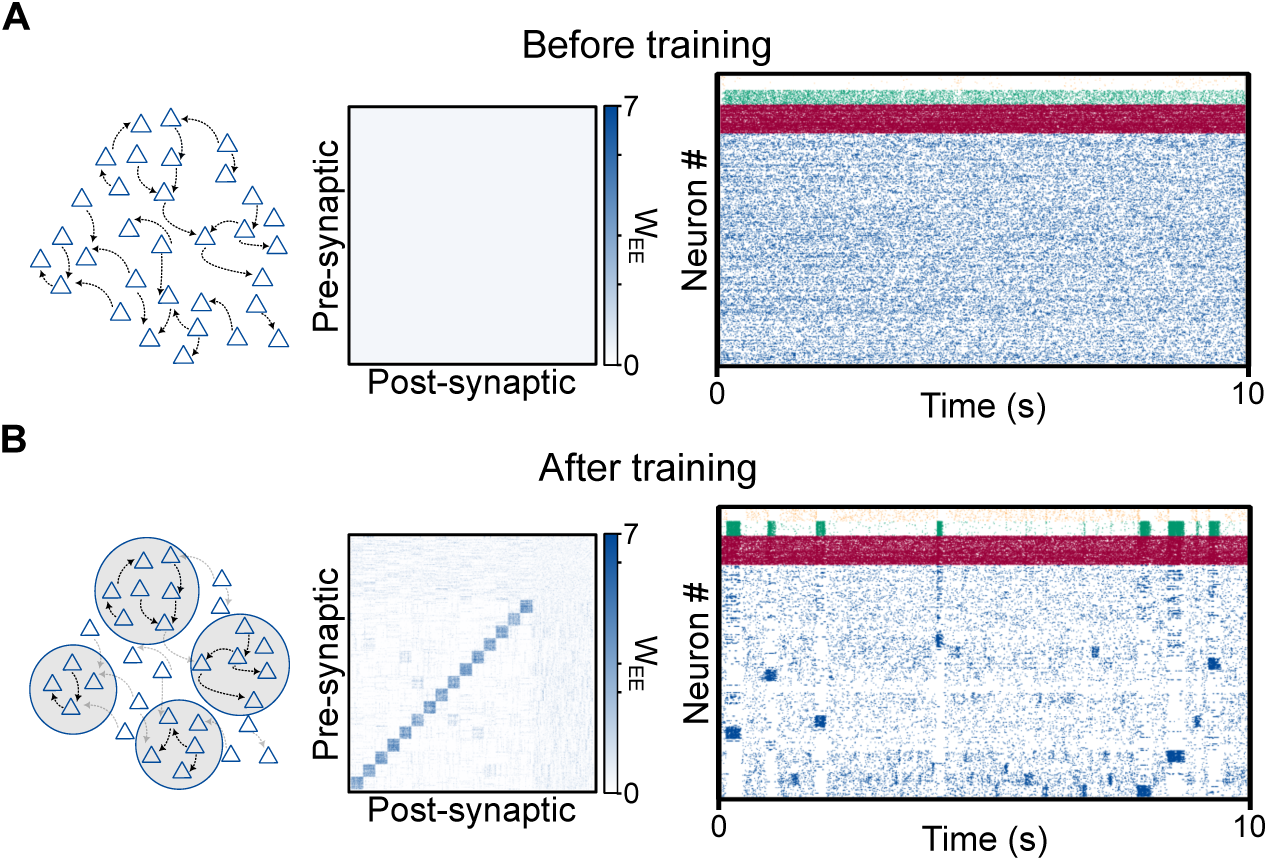
Training procedure equivalent to Figure 2 for IndNet. **A.** Before training procedure for IndNet showing: (left) schematic of homogeneous connectivity structure of excitatory connectivity, (middle) excitatory connectivity matrix, (right) raster plot showing excitatory (blue), PV (red), SST (green) and VIP (yellow) spikes over a time window of 10s. **B.** Same as **A.** but after training, showing the formation of assemblies thanks to the different plasticity mechanisms.

**Fig. S3.**
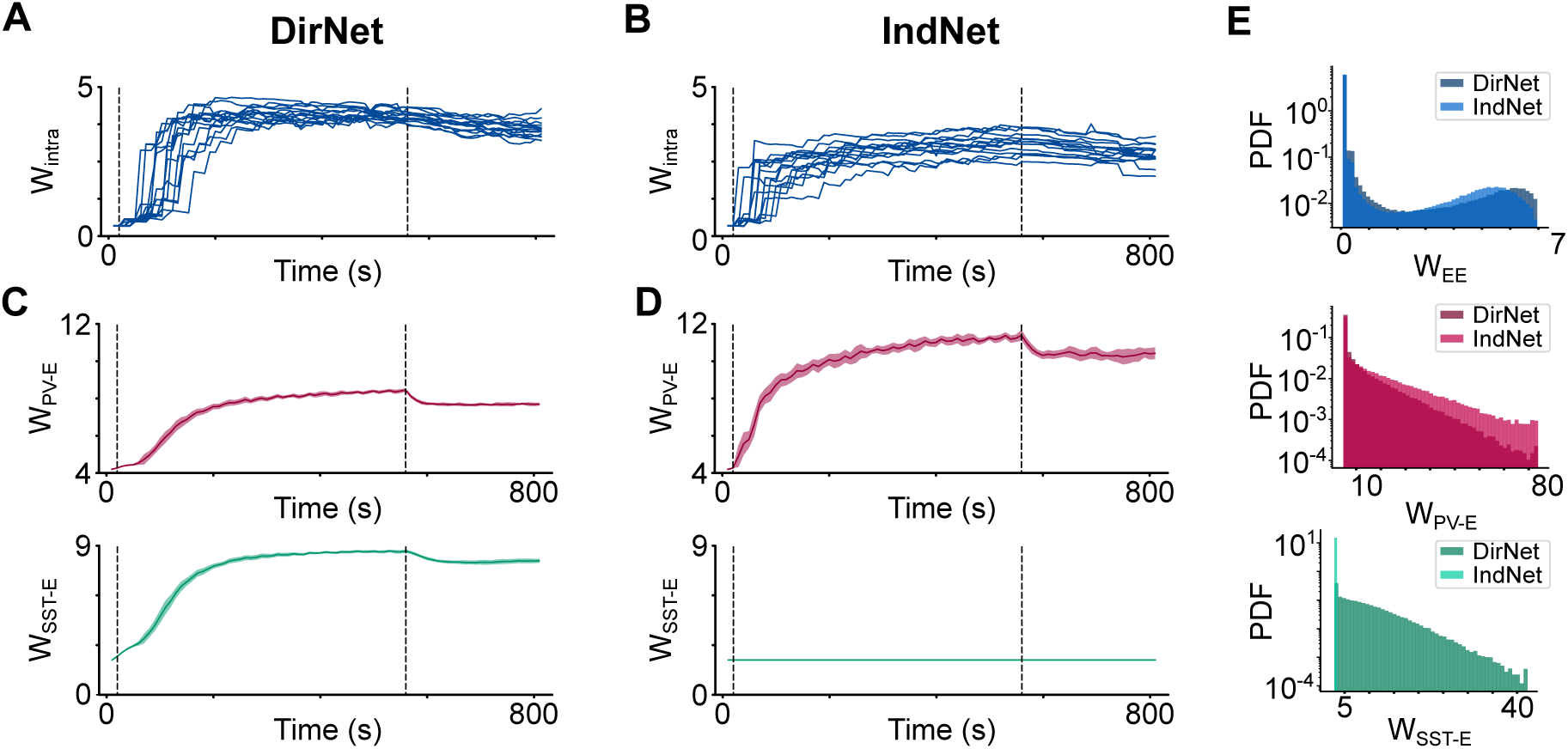
The imprinting of an assembly structure leads to metastable dynamics in both DirNet and IndNet. **A.** Synaptic weight evolution of the excitatory assemblies in DirNet. The dashed black lines indicate the training period in which excitatory assembly receives feedforward input. **B.** Same as **A.** for IndNet. **C.** Average synaptic weight evolution for PV-to-E (top) and SST-to-E (bottom) across trials of the inhibitory populations in DirNet. The dashed black lines indicate the training period in which excitatory assembly receives feedforward input. **D.** Same as **C.** for IndNet. **E.** Post-training excitatory (top), PV-to-E (middle) and SST-to-E (bottom) weight distributions for both DirNet (darker colors) and IndNet (lighter colors).

**Fig. S4.**
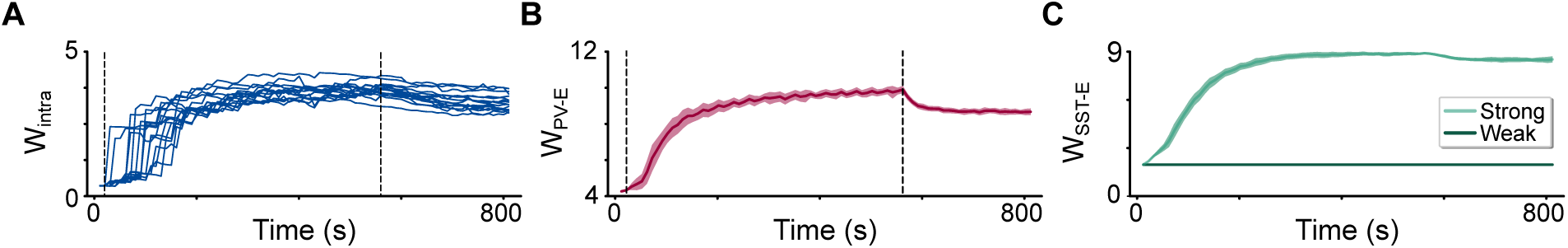
Training procedure in combined mixed network. **A.** Synaptic weight evolution of the excitatory assemblies. The dashed black lines indicate the training period in which excitatory assembly receives feedforward input. **B.** Average synaptic weight evolution for PV-to-excitatory across trials of the inhibitory populations. The dashed black lines indicate the training period in which excitatory assembly receives feedforward input. **C.** Same as **B.** for SST-to-excitatory connections. Light green represents SST-to-excitatory connections which become strong due to synaptic plasticity, whereas, dark green represents static connections which remain weak.

**Fig. S5.**
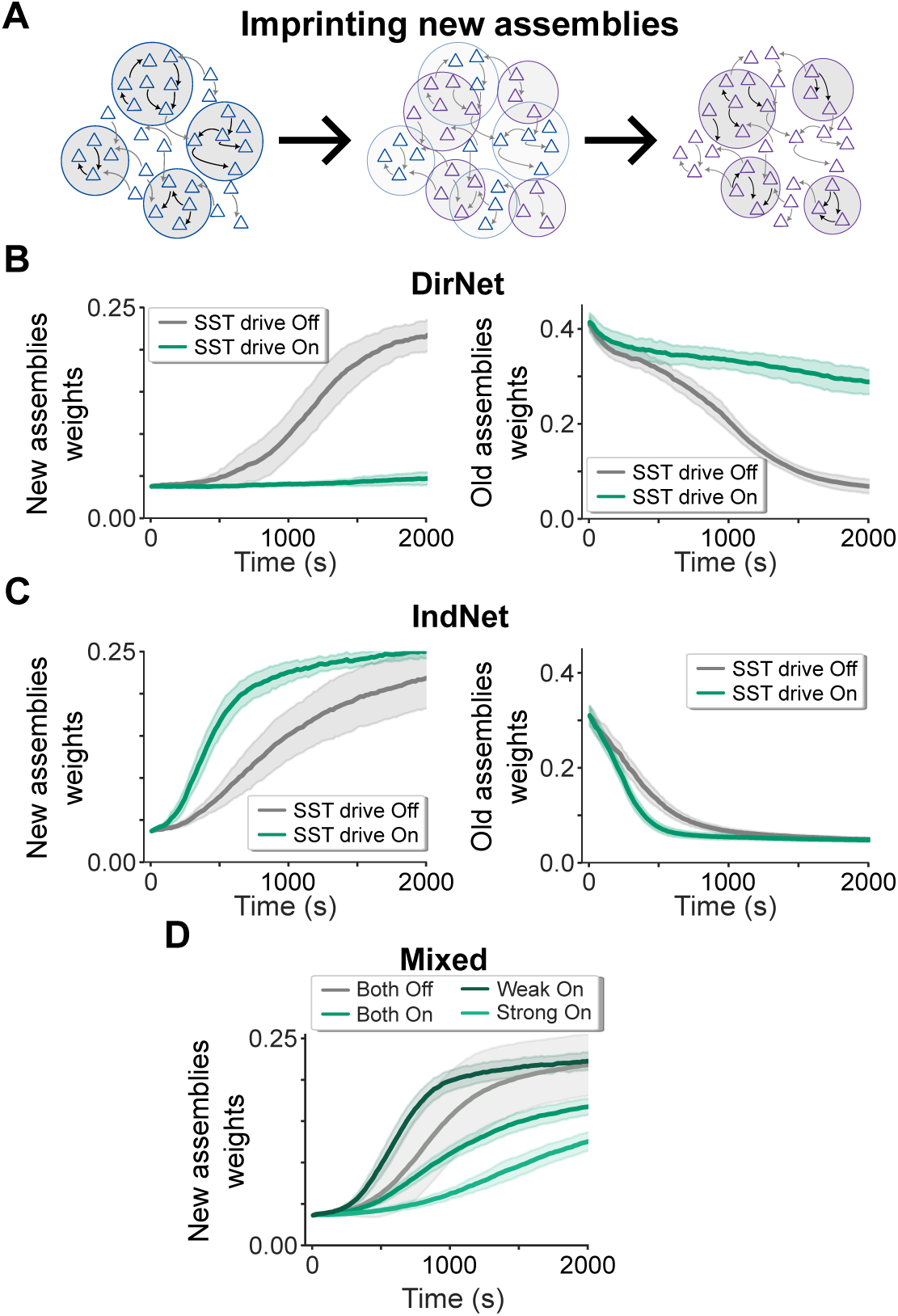
SST drive has opposite effects on the formation of new assembly structure in the two network configurations. **A.** Schematic of the reorganization of the original assembly structure (blue) into a new assembly structure (purple) generated by repartitioning the excitatory neurons into new subgroups. **B.** (Left) Evolution of the mean excitatory-to-excitatory synaptic weight within the new assemblies in DirNet with (green) and without (gray) SST drive. (Right) Evolution of the mean excitatory-to-excitatory synaptic weight within the original assemblies under the same conditions. **C.** Same as **B.** for IndNet. **D.** Weight evolution of the new excitatory assemblies in the mixed network without SST drive (gray) and while driving both SST subgroups (green), only the weak subgroup (dark green) or only the strong subgroup (light green).

**Fig. S6.**
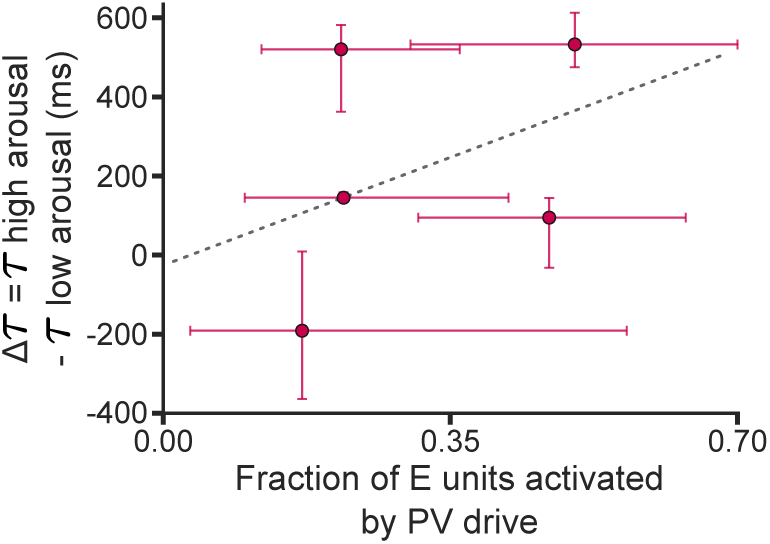
The fraction of PV-activated units does not predict the arousal-driven timescale change. Excitatory timescale difference between high and low arousal, Δ*τ* = *τ*_high_ *− τ*_low_, as a function of frac_up_, the fraction of E units activated by PV stimulation (*n* = 5 mice). Vertical error bars indicate bootstrap 95% confidence intervals (CIs) on Δ*τ*, and horizontal error bars indicate Wilson 95% CIs on frac_up_ (see Methods). The dashed line is a linear fit across mice (Pearson *r* = 0.40, *p* = 0.510; Spearman *ρ* = 0.60, *p* = 0.285).

**Table T1:**
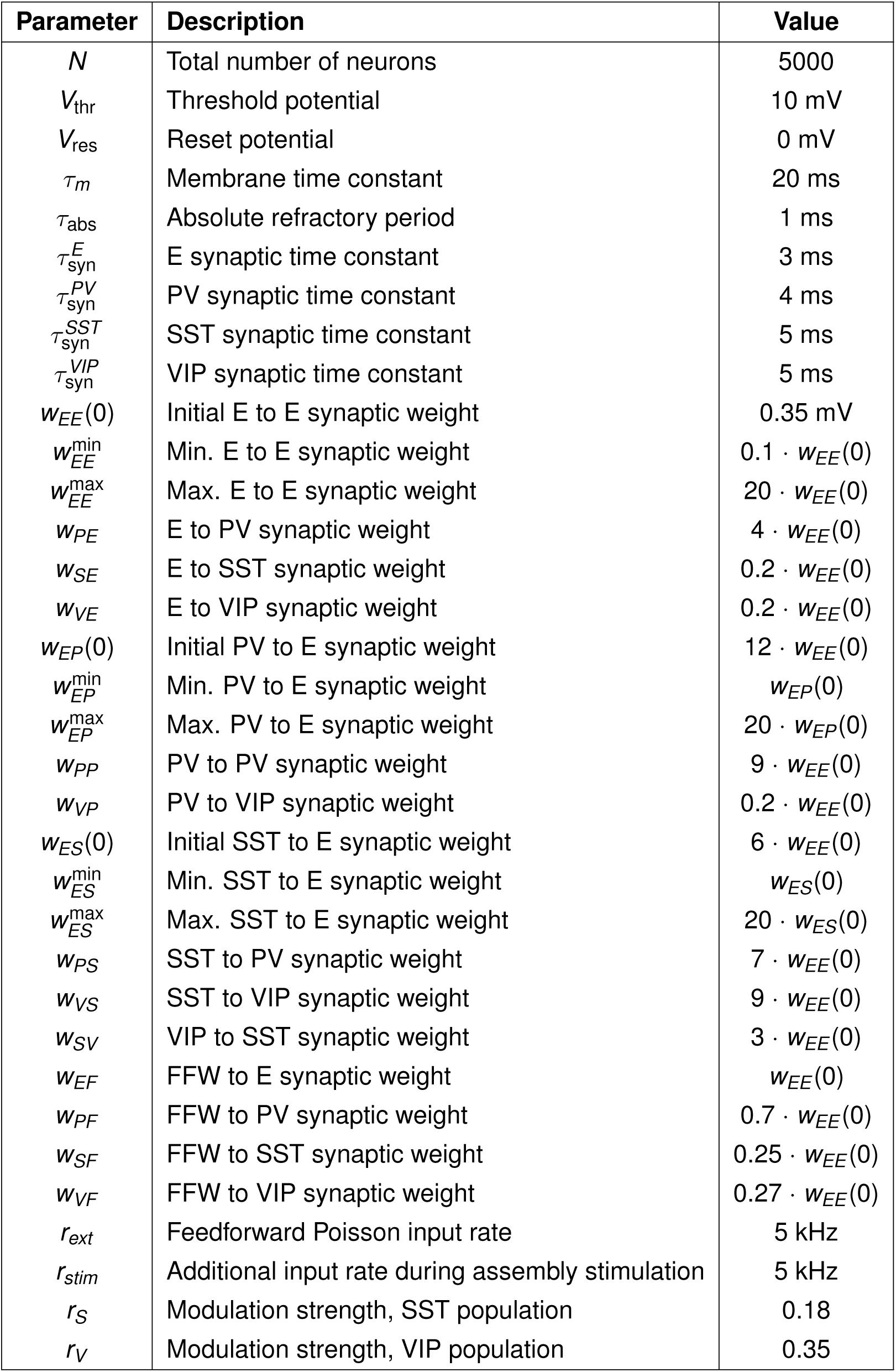
Neuron and network parameters.

| Parameter | Description | Value |
|---|---|---|
| *N* | Total number of neurons | 5000 |
| *V*<sub>thr</sub> | Threshold potential | 10 mV |
| *V*<sub>res</sub> | Reset potential | 0 mV |
| τ<sub>*m*</sub> | Membrane time constant | 20 ms |
| τ<sub>abs</sub> | Absolute refractory period | 1 ms |
| τ<sub>syn</sub><sup>*E*</sup> | E synaptic time constant | 3 ms |
| τ<sub>syn</sub><sup>*PV*</sup> | PV synaptic time constant | 4 ms |
| τ<sub>syn</sub><sup>*SST*</sup> | SST synaptic time constant | 5 ms |
| τ<sub>syn</sub><sup>*VIP*</sup> | VIP synaptic time constant | 5 ms |
| *w*<sub>*EE*</sub>(0) | Initial E to E synaptic weight | 0.35 mV |
| *w*<sub>*EE*</sub><sup>min</sup> | Min. E to E synaptic weight | 0.1 · *w*<sub>*EE*</sub>(0) |
| *w*<sub>*EE*</sub><sup>max</sup> | Max. E to E synaptic weight | 20 · *w*<sub>*EE*</sub>(0) |
| *w*<sub>*PE*</sub> | E to PV synaptic weight | 4 · *w*<sub>*EE*</sub>(0) |
| *w*<sub>*SE*</sub> | E to SST synaptic weight | 0.2 · *w*<sub>*EE*</sub>(0) |
| *w*<sub>*VE*</sub> | E to VIP synaptic weight | 0.2 · *w*<sub>*EE*</sub>(0) |
| *w*<sub>*EP*</sub>(0) | Initial PV to E synaptic weight | 12 · *w*<sub>*EE*</sub>(0) |
| *w*<sub>*EP*</sub><sup>min</sup> | Min. PV to E synaptic weight | *w*<sub>*EP*</sub>(0) |
| *w*<sub>*EP*</sub><sup>max</sup> | Max. PV to E synaptic weight | 20 · *w*<sub>*EP*</sub>(0) |
| *w*<sub>*PP*</sub> | PV to PV synaptic weight | 9 · *w*<sub>*EE*</sub>(0) |
| *w*<sub>*VP*</sub> | PV to VIP synaptic weight | 0.2 · *w*<sub>*EE*</sub>(0) |
| *w*<sub>*ES*</sub>(0) | Initial SST to E synaptic weight | 6 · *w*<sub>*EE*</sub>(0) |
| *w*<sub>*ES*</sub><sup>min</sup> | Min. SST to E synaptic weight | *w*<sub>*ES*</sub>(0) |
| *w*<sub>*ES*</sub><sup>max</sup> | Max. SST to E synaptic weight | 20 · *w*<sub>*ES*</sub>(0) |
| *w*<sub>*PS*</sub> | SST to PV synaptic weight | 7 · *w*<sub>*EE*</sub>(0) |
| *w*<sub>*VS*</sub> | SST to VIP synaptic weight | 9 · *w*<sub>*EE*</sub>(0) |
| *w*<sub>*SV*</sub> | VIP to SST synaptic weight | 3 · *w*<sub>*EE*</sub>(0) |
| *w*<sub>*EF*</sub> | FFW to E synaptic weight | *w*<sub>*EE*</sub>(0) |
| *w*<sub>*PF*</sub> | FFW to PV synaptic weight | 0.7 · *w*<sub>*EE*</sub>(0) |
| *w*<sub>*SF*</sub> | FFW to SST synaptic weight | 0.25 · *w*<sub>*EE*</sub>(0) |
| *w*<sub>*VF*</sub> | FFW to VIP synaptic weight | 0.27 · *w*<sub>*EE*</sub>(0) |
| *r*<sub>*ext*</sub> | Feedforward Poisson input rate | 5 kHz |
| *r*<sub>*stim*</sub> | Additional input rate during assembly stimulation | 5 kHz |
| *r*<sub>*S*</sub> | Modulation strength, SST population | 0.18 |
| *r*<sub>*V*</sub> | Modulation strength, VIP population | 0.35 |

**Table T2:** Parameters for synaptic plasticity.

| **Parameter** | **Description** | **Value** |
|---|---|---|
| *A*<sub>2</sub><sup>+</sup> | Pair potentiation amplitude | 5.0 · 10<sup>−10</sup> mV |
| *A*<sub>2</sub><sup>−</sup> | Pair depression amplitude | 7 · 10<sup>−3</sup> mV |
| *A*<sub>3</sub><sup>+</sup> | Triplet potentiation amplitude | 6.2 · 10<sup>−3</sup> mV |
| *A*<sub>3</sub><sup>−</sup> | Triplet depression amplitude | 2.3 · 10<sup>−4</sup> mV |
| *η*<sub>*PV*</sub> | iSTDP learning rate | 0.1 mV |
| *a*<sub>*pre*−*SST*</sub> | Asymmetric STDP pre-synaptic term | 0.81 mV |
| *a*<sub>*post*−*SST*</sub> | Asymmetric STDP post-synaptic term | 0.9 mV |
| *τ*<sub>+</sub> | Time constant for detector *r*<sub>1</sub> | 16.8 ms |
| *τ*<sub>*x*</sub> | Time constant for detector *r*<sub>2</sub> | 101.0 ms |
| *τ*<sub>−</sub> | Time constant for detector *o*<sub>1</sub> | 33.7 ms |
| *τ*<sub>*y*</sub> | Time constant for detector *o*<sub>2</sub> | 125 ms |
| *τ*<sub>*PV*</sub> | Time constant for iSTDP | 20 ms |
| *τ*<sub>*SST*</sub> | Time constant for asymmetric STDP | 30 ms |
| *ρ*<sub>0</sub> | Target firing rate | 3 Hz |
| Δ*t*<sub>*norm*</sub> | Normalization interval | 20 ms |

**Table T3:**
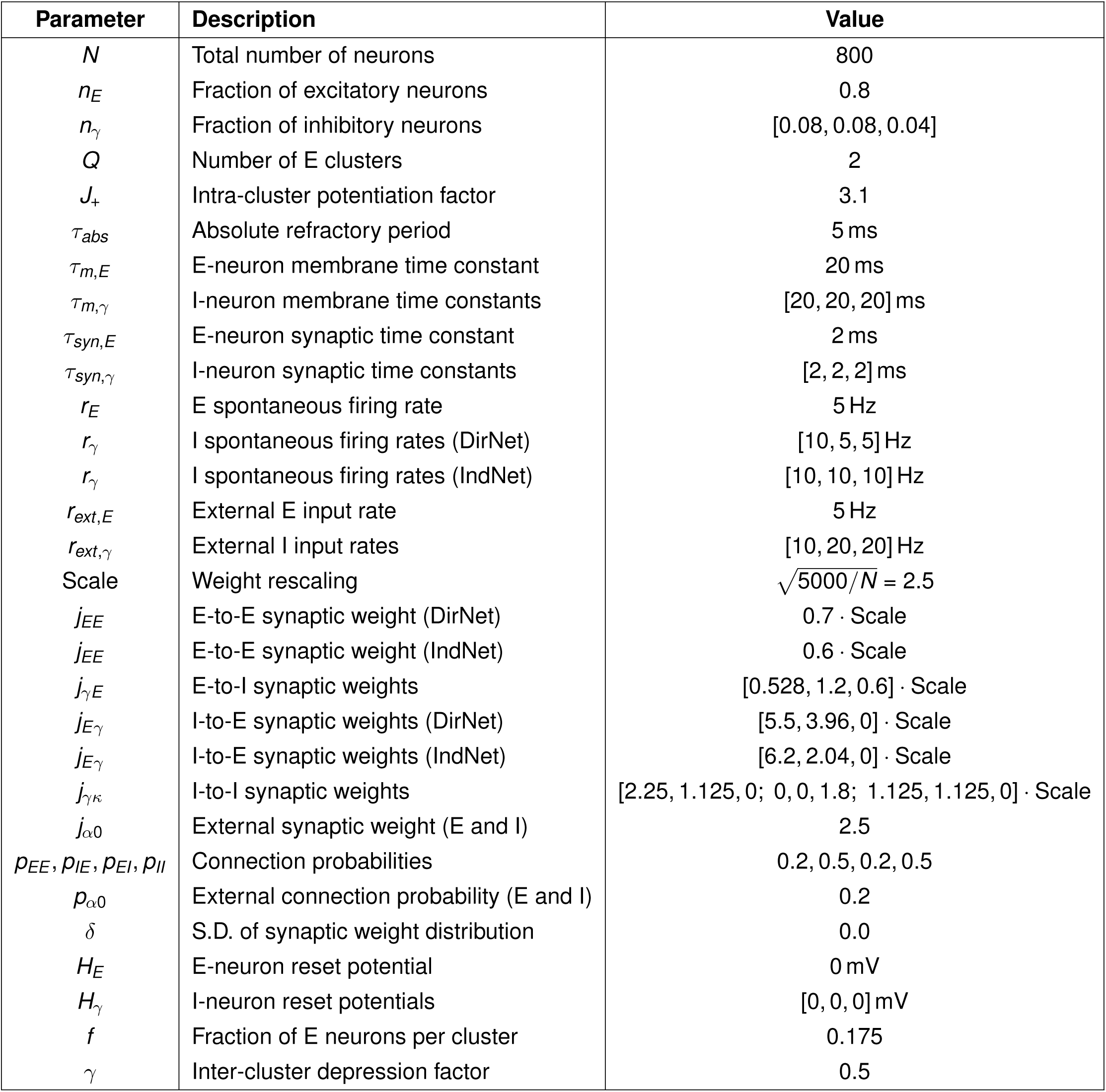
Parameters for mean-field approximation.

| Parameter | Description | Value |
|---|---|---|
| *N* | Total number of neurons | 800 |
| *n*<sub>*E*</sub> | Fraction of excitatory neurons | 0.8 |
| *n*<sub>γ</sub> | Fraction of inhibitory neurons | [0.08, 0.08, 0.04] |
| *Q* | Number of E clusters | 2 |
| *J*<sub>+</sub> | Intra-cluster potentiation factor | 3.1 |
| τ<sub>*abs*</sub> | Absolute refractory period | 5 ms |
| τ<sub>*m*,*E*</sub> | E-neuron membrane time constant | 20 ms |
| τ<sub>*m*,γ</sub> | I-neuron membrane time constants | [20, 20, 20] ms |
| τ<sub>*syn*,*E*</sub> | E-neuron synaptic time constant | 2 ms |
| τ<sub>*syn*,γ</sub> | I-neuron synaptic time constants | [2, 2, 2] ms |
| *r*<sub>*E*</sub> | E spontaneous firing rate | 5 Hz |
| *r*<sub>γ</sub> | I spontaneous firing rates (DirNet) | [10, 5, 5] Hz |
| *r*<sub>γ</sub> | I spontaneous firing rates (IndNet) | [10, 10, 10] Hz |
| *r*<sub>*ext*,*E*</sub> | External E input rate | 5 Hz |
| *r*<sub>*ext*,γ</sub> | External I input rates | [10, 20, 20] Hz |
| Scale | Weight rescaling | $\sqrt{5000/N}$ = 2.5 |
| *j*<sub>*EE*</sub> | E-to-E synaptic weight (DirNet) | 0.7 · Scale |
| *j*<sub>*EE*</sub> | E-to-E synaptic weight (IndNet) | 0.6 · Scale |
| *j*<sub>γ*E*</sub> | E-to-I synaptic weights | [0.528, 1.2, 0.6] · Scale |
| *j*<sub>*E*γ</sub> | I-to-E synaptic weights (DirNet) | [5.5, 3.96, 0] · Scale |
| *j*<sub>*E*γ</sub> | I-to-E synaptic weights (IndNet) | [6.2, 2.04, 0] · Scale |
| *j*<sub>γκ</sub> | I-to-I synaptic weights | [2.25, 1.125, 0; 0, 0, 1.8; 1.125, 1.125, 0] · Scale |
| *j*<sub>α0</sub> | External synaptic weight (E and I) | 2.5 |
| *p*<sub>*EE*</sub>, *p*<sub>*IE*</sub>, *p*<sub>*EI*</sub>, *p*<sub>*II*</sub> | Connection probabilities | 0.2, 0.5, 0.2, 0.5 |
| *p*<sub>α0</sub> | External connection probability (E and I) | 0.2 |
| δ | S.D. of synaptic weight distribution | 0.0 |
| *H*<sub>*E*</sub> | E-neuron reset potential | 0 mV |
| *H*<sub>γ</sub> | I-neuron reset potentials | [0, 0, 0] mV |
| *f* | Fraction of E neurons per cluster | 0.175 |
| γ | Inter-cluster depression factor | 0.5 |

